# Creating Red Light-Controlled Protein Dimerization Systems as Genetically Encoded Actuators with High Specificity

**DOI:** 10.1101/2020.07.09.174003

**Authors:** Zhimin Huang, Zengpeng Li, Xiao Zhang, Runze Dong, Shoukai Kang, Li Sun, Xiaonan Fu, David Vaisar, Kurumi Watanabe, Liangcai Gu

**Affiliations:** Department of Biochemistry and Institute for Protein Design, University of Washington, Seattle, WA 98195, United States; State Key Laboratory Breeding Base of Marine Genetic Resources, Third Institute of Oceanography, State Oceanic Administration, 184 University Road, Xiamen, 361005, P.R. China

**Author notes:** These authors contributed equally (Zhimin Huang, Zengpeng Li).

**Keywords:** light-induced dimerization, antibodies, protein engineering, biosensors, phage display

## Abstract

Protein dimerization systems that can be controlled by red light with increased tissue penetration depth are a highly needed optogenetic tool for clinical applications such as cell and gene therapies. However, existing red light-induced dimerization systems are all based on phytochrome photoreceptors and naturally occurring binding partners with complex structures and suboptimal *in vivo* performance, limiting mammalian applications. Here, we introduce an efficient, generalizable method (COMBINES-LID) for creating highly specific light-induced dimerization systems. Proof-of-principle was provided by creating <u>nano</u>body-based, <u>r</u>ed light-induced <u>d</u>imerization (nanoReD) systems comprising a truncated bacterial phytochrome sensory module using a mammalian endogenous chromophore, biliverdin, and light-form specific nanobodies. Selected nanoReD systems were biochemically characterized and exhibited low dark activity and high induction specificity for *in vivo* activation of gene expression. Overall, COMBINES-LID opens new opportunities for creating genetically encoded actuators for the optical manipulation of biological processes.

## Introduction

Since the invention of using light to control gene expression by Quail and coworkers in 2002,^[1]^ light-induced protein dimerization (LID) systems have been increasingly used to manipulate biological processes in living organisms.^[2,3]^ Similar to chemically induced dimerization (CID),^[4,5]^ LID systems can be genetically encoded to provide noninvasive control of molecular proximity, which is a fundamental regulator of gene regulation, cell signaling and metabolism, among other processes.^[6]^ For *in vivo* applications, LID offers spatiotemporal resolution that is unmatched by CID^[7]^ and, unlike chemical approaches, is not limited by toxicity, unintended effects of chemical inducers, or difficulties associated with drug delivery.

Different from single-component actuator systems such as microbial opsins,^[8]^ LID comprises two separate proteins or domains which serve as a sensor and an effector. The sensory function is initiated by i) light-induced chromophore isomerization or chromophore–protein bond formation triggering a conformational change of a chromophore-bound photosensory protein (hereafter named ‘*conformation switcher*’), or ii) photolytic release of a caged ligand or isomerization of a photoswitchable ligand that serves as a dimerization inducer.^[9]^ Naturally occurring conformation switchers widely exist in all kingdoms of life and many have been identified and characterized in the past three decades (Table S1). They have diverse structural and optical properties, offering flexible choices for *in vivo* applications. Many use widely shared metabolites from bacteria to humans as chromophores; for example, riboflavin-5’-phosphate bound to light-oxygen-voltage (LOV) sensing domains^[10]^ and biliverdin, a heme-derived linear tetrapyrrole found in bacterial phytochromes (BphPs).^[11]^

The effector function of LID is executed by a ‘*dimerization binder*’ which specifically binds to the conformation switcher in its light form—the state after a light-induced conformational change occurring to its thermally stable state in the dark, or the dark form (Scheme 1a). Many natural conformation switchers do not modulate protein–protein interaction and are not associated with any known dimerization binders; for example, in phytochromes, conformation switcher domains (known as a photosensory module) are typically fused with an enzymatic domain or module to allosterically regulate catalytic activity.^[12]^ Although a few natural dimerization binders have been identified (Table S1), they are limited by the basal activity in the dark (or dark activity) and other undesirable properties. It is yet difficult to design new dimerization binders with suitable specificity, sensitivity, and kinetics.

For deep-tissue applications in animals, LID is required to sense an optical input in the 650‒900 nm region, known as a tissue transparency window,^[13]^ because tissue absorbance, autofluorescence, and light scattering are minimized in this region.^[14,15]^ Phytochromes including plant and cyanobacterial phytochromes, BphPs, and cyanobacteriochromes use a class of linear tetrapyrroles, bilins, as chromophores that can sense low-energy optical signals in the far-red and near-infrared (NIR) range.^[16]^ Among those, BphPs use biliverdin, a ubiquitous endogenous bilin in mammalian cells,^[11]^ thus avoiding exogenous administration of the chromophore. However, so far only a natural dimerization binder, PpsR2, a ~50 kilodalton (kDa) transcriptional regulatory protein, was identified to bind to *Rhodopseudomonas palustris* BphP1 (*Rp*BphP1), a ~160 kDa homodimeric protein.^[17]^ PpsR2 was shown to interact with DNA and self-assemble as a dimer or higher oligomers,^[18]^ which can result in unwanted cellular outputs.^[19,20]^ Thus, a truncated form of PpsR2, Q-PAS1, was generated to eliminate DNA binding while maintain binding to *Rp*BphP1.^[20]^ However, both PpsR2 and Q-PAS1 LIDs exhibited a significant dark activity,^[20,21]^ likely due to the nonspecific binding to the dark form of *Rp*BphP1.

**Scheme 1.**
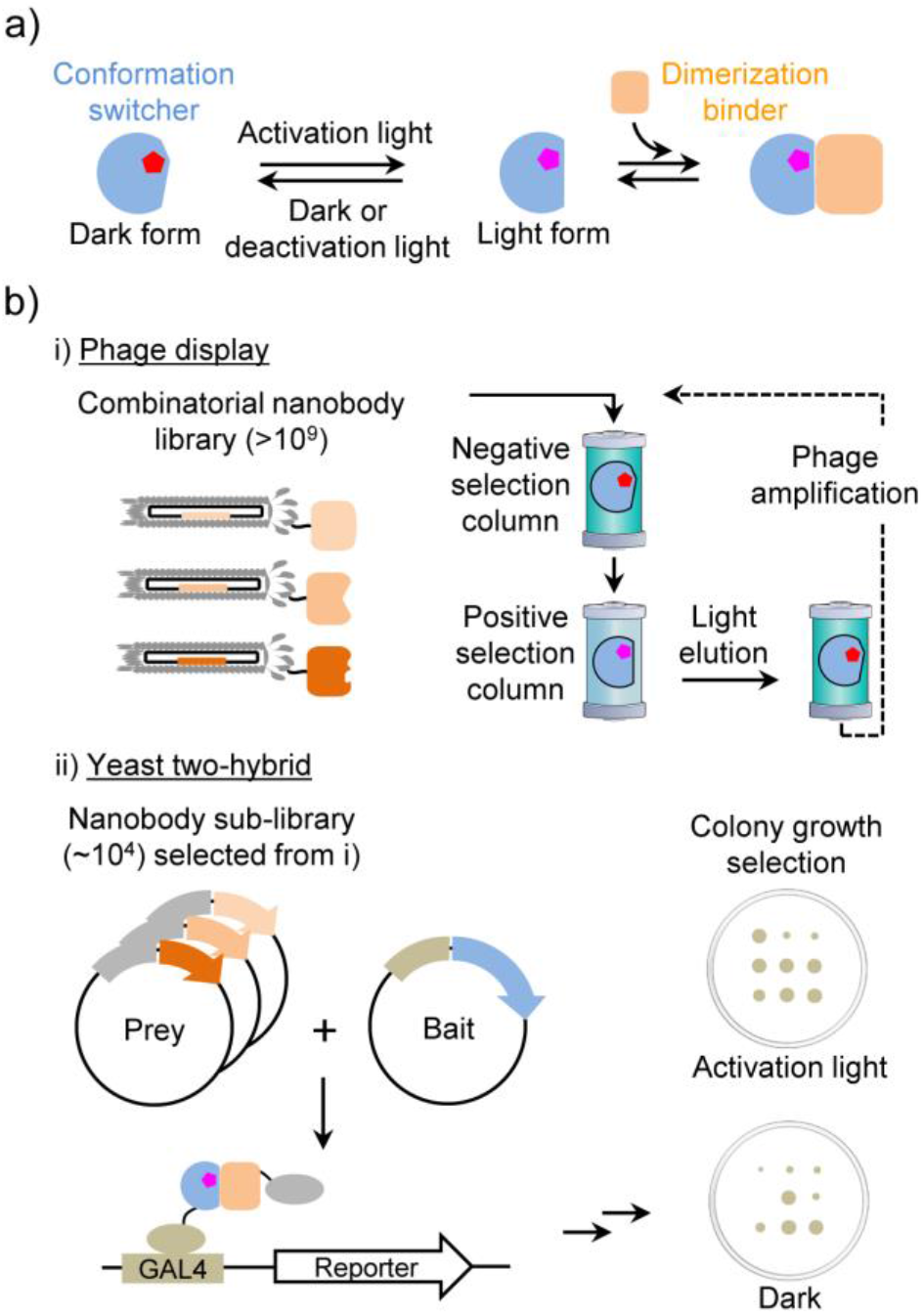
a) LID mechanism. b) Principle of the COMBINES-LID method.

Here we aimed to create LID systems with decreased protein sizes and minimized dark activity by *de novo* engineering of small dimerization binders to a photosensory module excised from a full-length BphP.^[22,23]^ So far, creating light form-specific binders was mainly achieved by *in vitro* selection due to the feasibility to manipulate light sensitive proteins in a screening assay. Initial successes have been reported using phage display to screen computationally designed binders for a LOV2 domain^[24]^ and a random surface mutation library of an albumin-binding domain targeting the LOV2 and a photoactive yellow protein.^[25]^ Inspired by these works, we sought to establish a robust, general method by coupling *in vitro* and cell-based screening to create new LID systems for mammalian applications. To facilitate implementation by other labs, we chose to screen one of the mostly used small binders, single-domain antibody (or nanobody), a 12-15 kDa functional antibody domain with a universal scaffold and three variable complementarity-determining regions (CDRs).^[26]^

## Results and Discussion

We devised a two-step screening method, <u>comb</u>inatorial <u>bin</u>ders-<u>e</u>nabled <u>s</u>election of LID (COMBINES-LID), which involves phage display to enrich binders that only bind to the BphP light form and then yeast two-hybrid (Y2H) screening of the enriched sub-library to select for *in vivo* activity (Scheme 1b). A high-quality synthetic combinatorial nanobody library generated in our previous work^[27]^ was used. These nanobodies have an optimized scaffold^[28]^ and rationally randomized CDRs with an estimated sequence diversity of 1.23 to 7.14 × 10^9^.

To simplify the structure of the conformation switcher, the photosensory module of *Deinococcus radiodurans* phytochrome (*Dr*BphP) was chosen. Its light and dark forms can be photoconverted by activating far-red (e.g., 654 nm) and deactivating NIR (e.g., 775 nm) illuminations, respectively, and then stably maintained in a screening assay. The photoswitching efficiency is close to that of the full-length *Dr*BphP.^[29,30]^ By contrast, the excised module of *Rp*BphP1 showed impaired photoconversion.^[17]^ The tridomain module (hereafter named *Dr*BphP for simplicity) comprising a Per-ARNT-Sim (PAS), a cGMP phosphodiesterase-adenylate cyclase-FhlA (GAF), and a phytochrome-specific (PHY) domains was expressed as a ~60 kDa fusion bearing a C-terminal AviTag and HisTag, incubated with biliverdin, purified, and biotinylated (Figure S1) to serve as a bait for phage display. The photoconversion of the purified protein was confirmed by measuring the spectra of the light and dark forms (Figure S2).

We hypothesized that specific and reversible dimerization binders are critical for the *in vivo* performance of LID, such as a low dark activity. To enhance selection efficiency, we used column chromatography to continuously separate phage-displayed nanobodies between the stationary and mobile phases as they passed through a column. Binding specificity was selected by loading the library onto two connected transparent columns, the first (negative selection) preloaded with biotinylated *Dr*BphP in the dark form and the second (positive selection) with the light form (Figure S3a). Thus, nanobodies captured in the second column should have zero or very low affinity to the dark form. Next, reversible binders were collected by eluting only dissociated nanobodies from the second column after switching the light to dark form by the 775-nm illumination. After four-round phage binding and elution with gradually decreased illumination times (Figure S3b), the final eluent was estimated to contain ~90% light-eluted clones (Table S2) with the sequence diversity of ~10^4^.

*In vitro* selected nanobodies were subcloned into a Y2H sub-library for the cell-based screening of cytoplasmic expression and binding specificity. Although some nanobodies can be functionally expressed as intracellular binders,^[28]^ many are unstable leading to loss of function due to the inability of disulfide bond formation in the reducing cytoplasmic environment. Y2H was selected for the sub-library screening due to its suitable throughput and cost-effectiveness. Y2HGold cells were co-transformed with plasmids carrying genes of *Dr*BphP and nanobodies and selected on SD/-Ade/-His/-Leu/-Trp agar plates under the 654-nm illumination. ~2,000 fully grown colonies were picked, inoculated into 1-mL SD/-Leu/-Trp medium, and replica spotted onto the agar plates to compare colony growth under the illumination and in the dark (Figure S4). Five candidates grew only under the illumination (Figure 1a) and with diverse CDR sequences (Table S3) were selected for further characterization.

**Figure 1.**
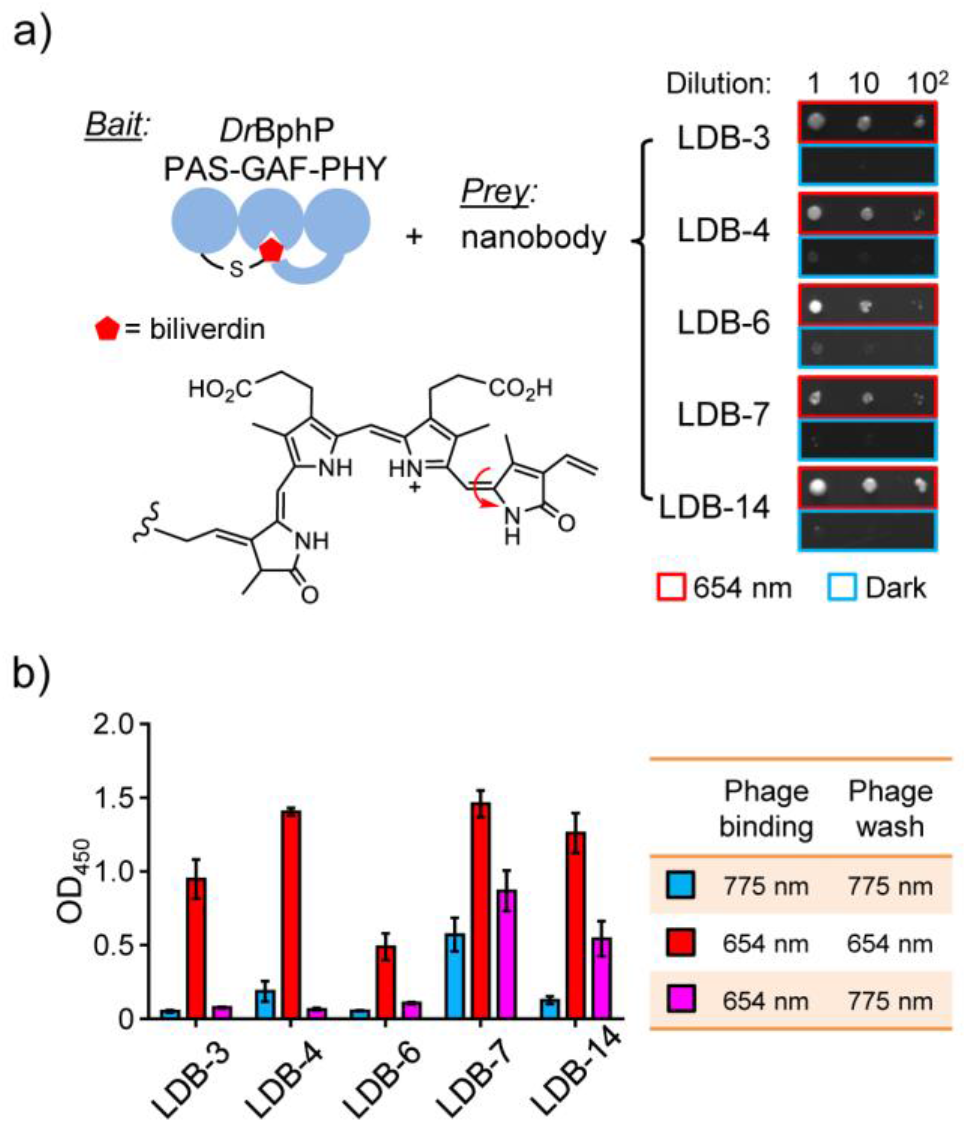
Y2H and single phage ELISA analyses of dimerization binder candidates. a) Y2H assay with the biliverdin-bound *Dr*BphP photosensory module as a bait and nanobodies as preys. A serial dilution of Y2HGold cells resuspended in 0.9% NaCl were spotted on SD/-Ade/-His/-Leu/-Trp plates and grown under the 654-nm illumination (0.03 mW/cm^2^) or in the dark. A representative result from three independent experiments is shown on the right. b) ELISA analysis of nanobody binding specificity and reversibility. Phage-displayed nanobodies were bound to *Dr*BphP immobilized in microtiter plates, which were illuminated with the 654-nm (0.3 mW/cm^2^) or 775-nm (0.2 mW/cm^2^) lights during the binding and wash steps. Data represent mean values of 3 measurements; error bars, standard deviation.

To confirm the binding specificity and reversibility, we assayed selected nanobodies by single phage enzyme-linked immunosorbent assay (ELISA). Phage displayed-nanobodies were first bound to the dark and light forms of biotinylated *Dr*BphP immobilized in streptavidin-coated microtiter plates. To maintain the dark or light form, or to convert the light to dark form, the plates were under 654- and/or 775-nm illumination during the phage binding and wash steps. As expected, all candidates showed light-form binding specificity with non-detectable (LDB-3 and LDB-6) to relatively low (LDB-4, LDB-7, and LDB-14) binding to the dark form (Figure 1b). Bound nanobodies were almost completely (LDB-3, LDB-4, and LDB-6) or partially (LDB-7 and LDB-14) washed off after converting the light to dark form.

To determine whether nanobody candidates are suitable for mammalian applications, we tested their expression in human embryonic kidney 293T (HEK293T) cells. It is known that the same protein–protein interaction (PPI) found in yeast might not be detected in mammalian cells due to protein expression or stability issues; for example, a recent comparison of PPI assays in different hosts found that only half of human PPIs detected in yeast were also seen in HEK293T, and vice versa.^[31]^ Thus, we were interested to know the success rate of nanobodies selected by COMBINES-LID that can be functionally expressed in mammalian cells.

To compare *in vivo* activity, we assayed proteins by mammalian two-hybrid (M2H)^[32]^ under a standardized condition. Specifically, *Dr*BphP was fused with an N-terminal GAL4 DNA binding domain (BD) and nanobodies with a C-terminal p65 transcriptional activation domain (AD) to control the expression of a firefly luciferase reporter (Figure 2a). After transient co-transfection with the BD and AD, and reporter plasmids, cells were cultured for 24 hours in the dark to express the *Dr*BphP and nanobodies, and then maintained in the dark or under the 654-nm illumination for another 24 hours to compare the luciferase expression. Dark activity and specificity were analyzed by comparing the dark expression with a negative control (i.e., only *Dr*BphP was expressed) and the light-induced expression with the dark expression, respectively. All candidates showed low dark activities, and the dark expression levels were close to the control (Figure 2b). LDB-3 and LDB-14 showed the high specificity; their light-induced expression levels were increased by ~19.5 and ~19.1 folds, respectively. The other three candidates did not show obvious light activation. To understand their loss of the expected activity, we investigated protein stability in HEK293T. Specifically, all nanobodies bearing a C-terminal SNAP-tag were expressed, fluorescently labelled, and analyzed by sodium dodecyl sulfate gel electrophoresis (SDS-PAGE) to detect full-length proteins and degraded forms. All nanobodies were found with degraded fragments; however, compared with LDB-3 and LDB-14, the other three nanobodies showed drastically decreased levels of full-length proteins, suggesting that they might not be stable in the host cells (Figure S5).

**Figure 2.**
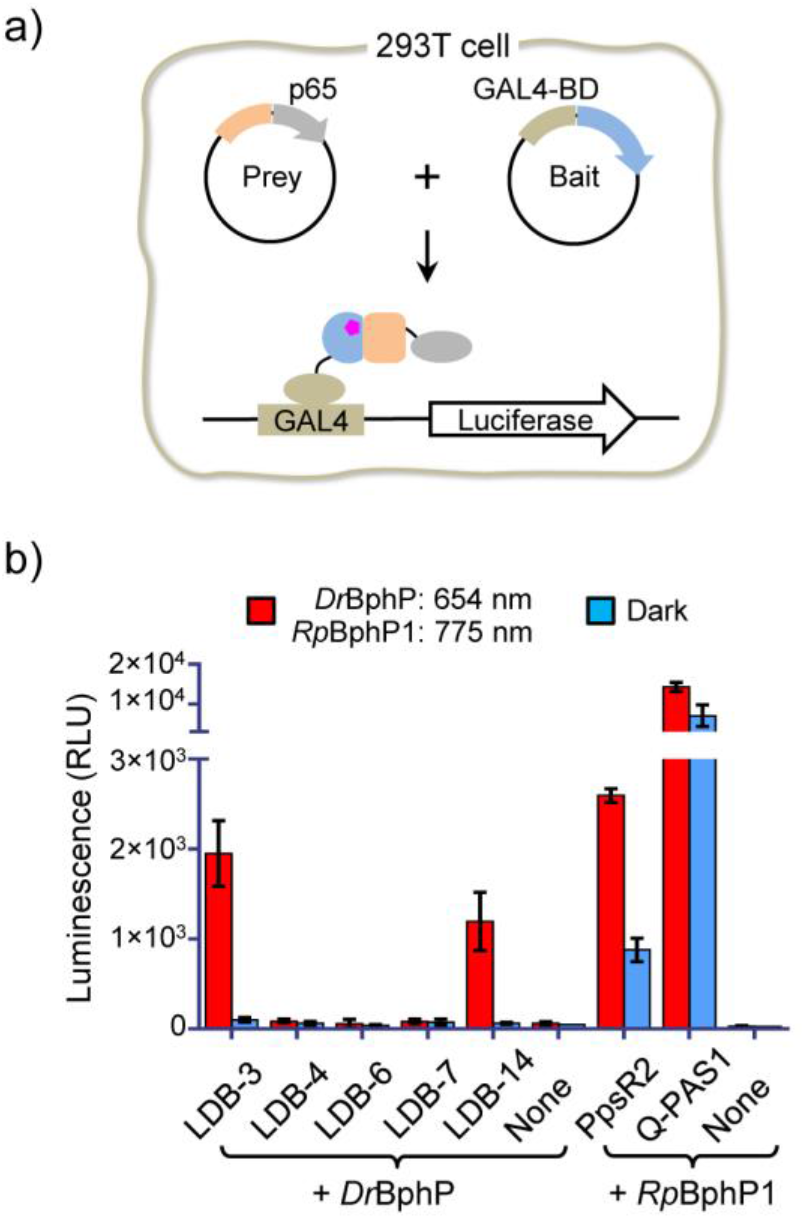
Nanobody specificity validation in mammalian cells. a) Schematic of the M2H assay. *Dr*BphP and nanobody genes were inserted into the bait and prey plasmids, respectively. b) Specificity comparison of LID systems. HEK293T cells were transiently co-transfected with the bait, prey, and GAL4UAS-luciferase reporter plasmids (~0.25 μg each) in a 0.5 mL culture. None, the negative control transfected with only the bait and the luciferase reporter plasmids. Cells were cultured under the illumination (654 nm (0.2 mW/cm^2^) or 775 nm (0.2 mW/cm^2^)) or in the dark for 24 hours before measuring luciferase levels. Different from *Dr*BphP, *Rp*BphP1 is required to be converted to the light form by a NIR (e.g., 775-nm) light. Data represent mean values of 3 measurements; error bars, standard deviation.

We compared the specificity of the nanobody-based LIDs with the *Rp*BphP1–PpsR2 and *Rp*BphP1–Q-PAS1 systems by M2H. The *Rp*BphP1-based LIDs showed light-enhanced expression (Figure 2b). However, due to the high dark activity, light-induced expression was only increased by ~2.95 and ~2.02 folds. Our observed specificity of the *Rp*BphP1 systems is much lower than originally reported,^[19,20]^ but is close to that recently achieved in *Escherichia coli*.^[33]^ As suggested by a mathematical model of LID,^[34]^ the specificity is determined by not only relative binding affinities of dimerization binders to dark and light forms of a conformation switcher, but also by their effective cellular concentrations, which unfortunately are difficult to measure in our experiment. Nevertheless, our comparison with the same transfection and culture condition suggests that LDB-3 and LDB-14 offer significantly enhanced dimerization specificity.

We biochemically assessed LDB-3 and LDB-14 to understand their binding mechanisms. We first sought to detect light-induced *Dr*BphP-nanobody complexes using analytical size-exclusion chromatography (SEC). LDB-3 and LDB-14 were bacterially expressed and purified with yields of ~2-3 milligrams per liter of culture. SEC data showed that LDB-3 was a mixture of the monomer and dimer and LDB-14 mainly the monomer and both nanobodies dimerized at increased concentrations (Figure S6a). Consistent with a previous report,^30^ both the light and dark forms of *Dr*BphP were eluted mainly as homodimers (Figure S6b). After *Dr*BphP was illuminated and then incubated with LDB-3 or LDB-14 in the dark, split SEC peaks of *Dr*BphP were observed only for the light form (Figure 3a), implying complex formation. Complexes were confirmed by SDS-PAGE detection of coeluted *Dr*BphP and nanobodies (Figure 3b). Interestingly, gel analysis revealed that LDB-3 was coeluted with *Dr*BphP later than LDB-14, suggesting that they might have different binding configurations or affinities. Consistent with the single phage ELISA result (Figure 1b), LDB-14 appeared to weakly interact with the dark form since a faint nanobody band was detected in the dark-form *Dr*BphP factions.

**Figure 3.**
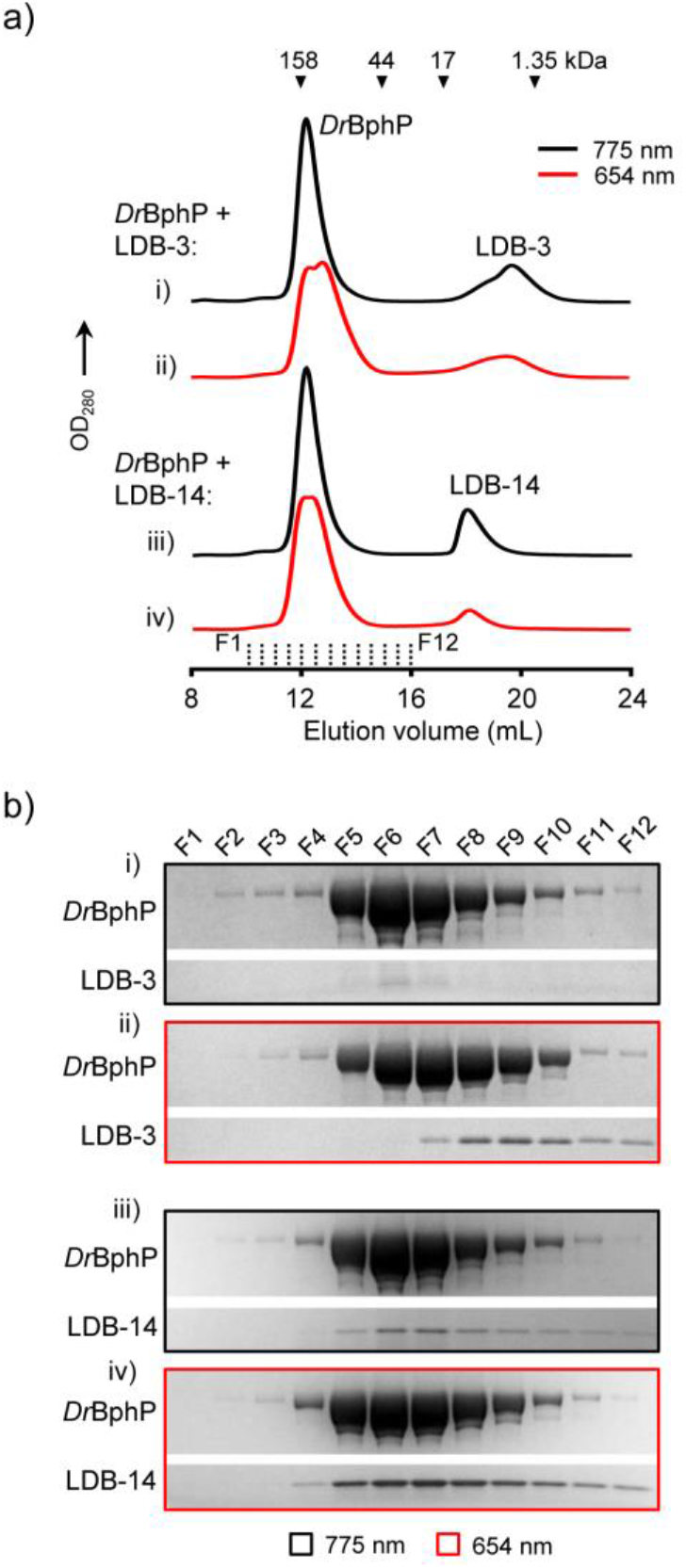
Detection of light-induced *Dr*BphP-nanobody complexes. a) Analytical SEC. ~6 μM (final concentration) *Dr*BphP after the 654-nm (0.2 mW/cm^2^) or 775-nm (0.8 mW/cm^2^) illumination for 5 min were incubated with ~5 μM (final concentration) LDB-3 or LDB-14 in the dark. 500 uL mixtures were loaded onto a Superdex 200 Increase 10/300 GL column pre-equilibrated with 1× PBS buffer and eluted at 0.75 mL/min at 4°C. Elution volumes of protein standards are marked by triangles. b) SDS-PAGE analysis of the fractions (500 μL each) collected in the SEC (marked by dash lines in a)) and concentrated by trichloroacetic acid precipitation. Only gel regions showing *Dr*BphP and nanobody bands are shown.

We next studied the thermodynamics of nanobody binding by isothermal titration calorimetry (ITC). Binding data obtained by titrating LDB-3 or LDB-14 into a photoconverted light-form sample were fitted using a one-site model (*R*^2^ = ~0.99) to give apparent dissociation constants (*K*D^app^s) of 1.01 and 0.47 μM, respectively (Figure 4 and Table S4). The binding site number of *Dr*BphP was calculated to be ~0.6, consistent with a previous small-angle X-ray scattering result that only 64% of the protein molecules in the photoconverted sample adopted the light-state conformation.^30^ Unexpectedly, the thermograph of LDB-3 titration into the dark form showed significant heat exchange: a clear transition from heat release to absorption when more LDB-3 was added, which was not observed for LDB-3 titration into the light form (Figure 4a). This transition suggests that the exothermic binding of LDB-3 to the dark form might be coupled to an endothermic process, which is likely to be the LDB-3 dimer dissociation (Figure S7). In other words, LDB-3 binding to the dark form might be inhibited by LDB-3 dimerization, thus providing a mechanism to explain the observed low dark activity of LDB-3 (refer to the thermodynamic simulation in Supplementary Note). LDB-14 was found to only bind to a small fraction (~15%) of *Dr*BphP in the dark-form sample but did not generate measurable heat of binding with the major population (Figure 4b). The minor fraction was likely the protein switched to the light form during our assay.

**Figure 4.**
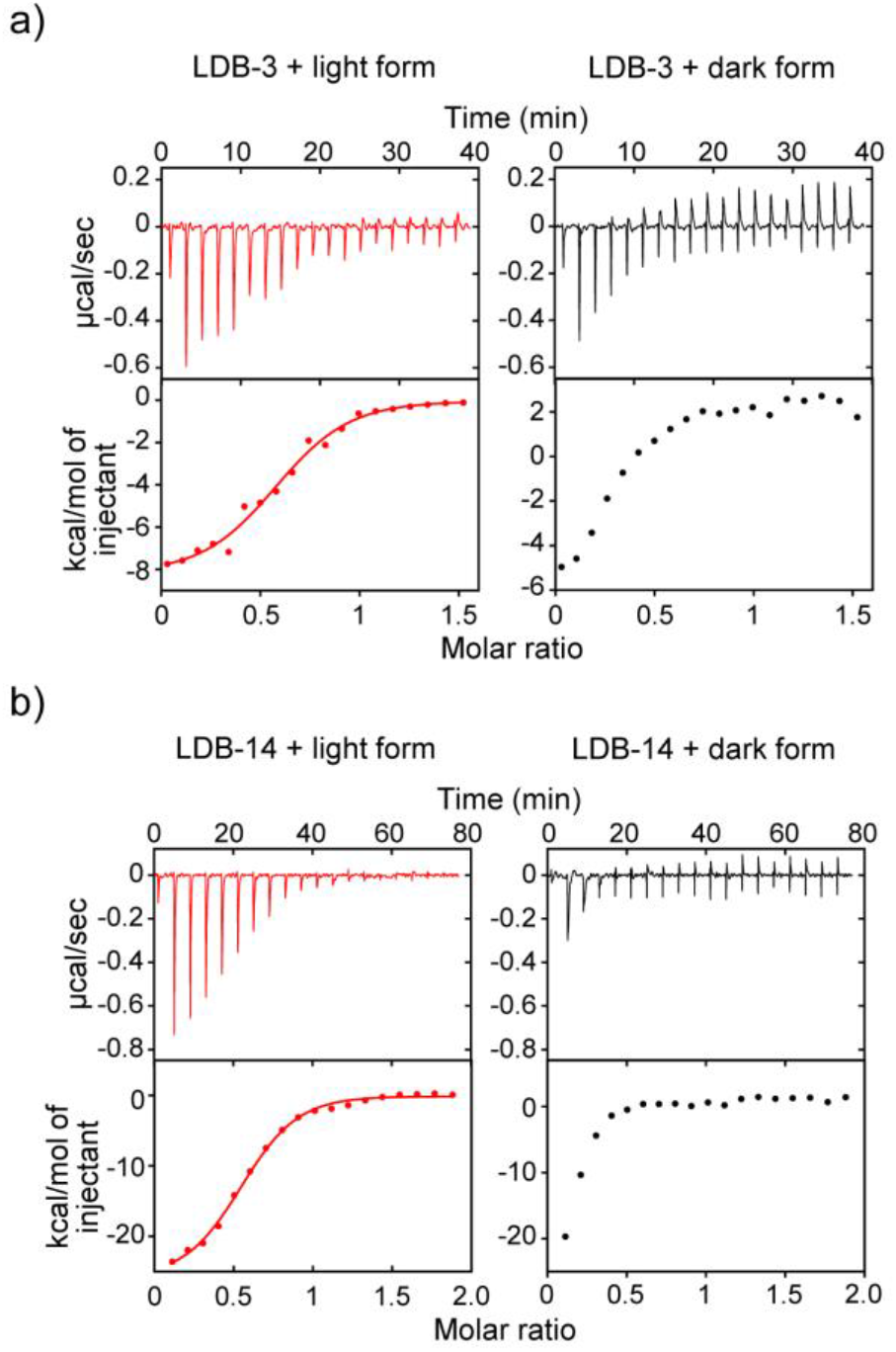
ITC thermographs of the nanobody binding. a) 80 μM LDB-3 and b) 50 μM LDB-14 were titrated into 10 μM and 5 μM DrBphP, respectively. The light and dark forms were converted by the 654-nm (0.2 mW/cm^2^) and 775-nm (0.2 mW/cm^2^) lights, respectively. The raw data (top) and the integration of heats (bottom) for each titration are shown.

The binding kinetics of LDB-3 and LDB-14 were measured by Bio-Layer Interferometry (BLI). The assay was performed by incubating the light or dark form with nanobodies immobilized on streptavidin biosensors (refer to Supplementary Methods). The result revealed that, compared with LDB-14, LDB-3 has a weaker binding affinity to the light form mainly due to a ~4.9-fold faster dissociation from the *Dr*BphP (*k*_off_ = ~18.5 × 10^-2^s^-1^) (Figure S8 and Table S5). Theoretically, the fast dissociation could offer higher temporal resolution for the reversible control of LID.^34^ The BLI analysis of the dark form binding was not straightforward because the white light signal used by this optical technique might partially convert the biosensor-bound *Dr*BphP to the light form.

Since *Dr*BphP photoconversion and nanobody binding might reciprocally affect each other, we asked whether the nanobody binding can slow the photoconversion to the dark state. To test this, we illuminated *Dr*BphP with different exposure times and light intensities and measured the percentage of the dark state in the protein by the ratio of absorption at 750 nm (A_750_) to 700 nm (A_700_) (Figure S9a). Compared with unbound *Dr*BphP, the nanobody-bound *Dr*BphP had decreased photoconversion rates (Figure S9b), suggesting that the nanobody binding can stabilize the light state. At a relatively low light intensity (0.05 mW/cm^2^), the LDB-3-bound *Dr*BphP had a slightly faster relaxation to the dark state than the LDB-14-bound *Dr*BphP, likely due to the faster dissociation of the *Dr*BphP– LDB-3 complex. Together, our biochemical data support the high specificity and reversibility of the nanobody-based LID systems.

To develop *in vivo* applications, we focused on light-induced gene expression. We first determined the time-course response of light-induced activation of luciferase expression in HEK293T cells. Under the same culture and transfection condition, luciferase levels with or without 654-nm illumination were measured at seven time points up to 72 hours. For both LDB-3 and LDB-14, luciferase levels after illumination reached half maximum and maximum at 12 and 24 hours, respectively (Figure S10). The half-maximum and maximum luciferase levels in cells expressing LDB-3 were ~2.5-fold higher than those of cells expressing LDB-14, which seems to be correlated with *in vivo* nanobody stability (Figure S5).

The luciferase assay required releasing the protein by cell lysis, so we also measured *in situ* green fluorescent protein (GFP) expression by fluorescence imaging. Specifically, HEK293T cells were transiently co-transfected with LID genes to control the transcription of a chromosomally integrated GFP gene. Imaging analysis showed zero to very low GFP expression in cells kept in the dark (Figure 5a). In LDB-3- and LDB-14-expressing cells, light-induced GFP levels were close and, compared with the expression in the dark, increased by ~44 and ~39 folds, respectively (Figure 5b).

**Figure 5.**
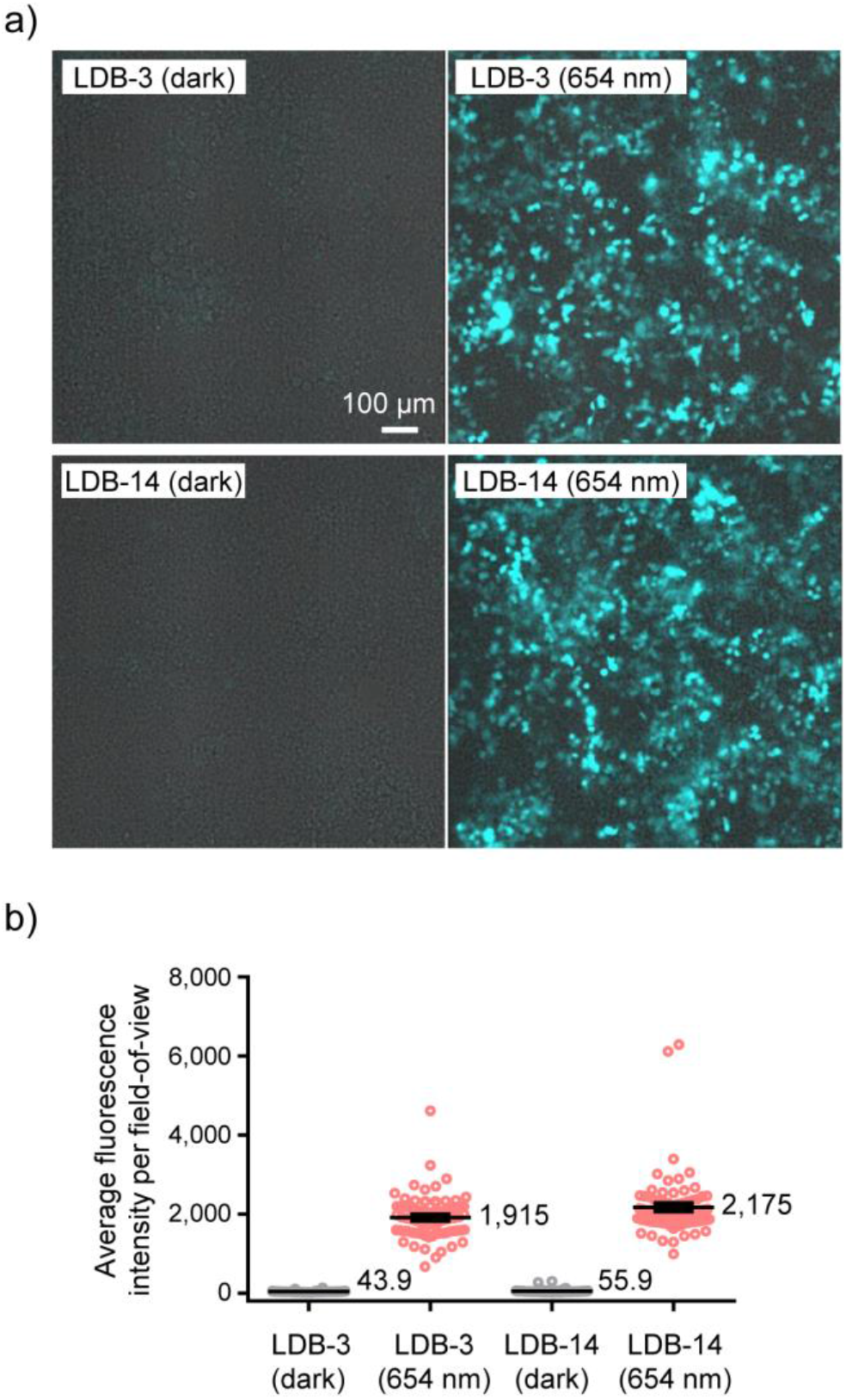
Red light-induced expression of GFP. a) Representative overlaid brightfield and fluorescence images of GFP-expressing HEK293T cells. Cells were transduced with a lentiviral GFP expression vector and then co-transfected with the LID plasmids after 654-nm (0.2 mW/cm^2^) illumination or in the dark for 48 hours. b) Comparison of GFP fluorescence intensities in fields-of-view (FOVs). Data represent mean values of 78 FOVs; error bars, standard error of the mean (SEM).

Finally, to demonstrate the gene activation in living animals, we subcutaneously injected the LDB-3/*Dr*BphP-controlled luciferase-expressing HEK293T cells into nude mice, remotely activated the transcription by the 654-nm light, and quantified luciferase expression by bioluminescence imaging. As expected, mice kept in the dark were detected with low bioluminescence signals and those received the 24-hour illumination had a ~25-fold average increase of bioluminescence (Figure 6), which is consistent with the data obtained with the cultured cells (Figure S10). Considering cells had been exposed to light during the subcutaneous injection, the dark activity is expected to be even lower if light exposure can be completely avoided.

**Figure 6.**
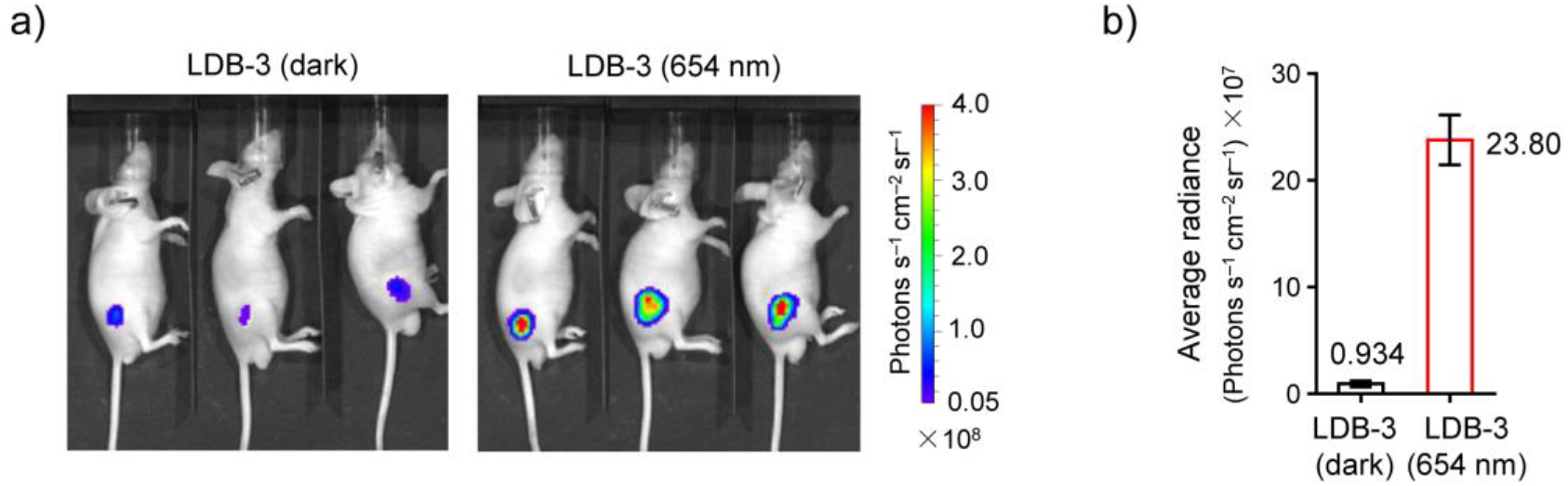
Red light-induced transcription activation in mice. a) Bioluminescence imaging of nude mice which were subcutaneously injected with HEK293T cells co-transfected with LID and the luciferase reporter plasmids. The luciferase expression was measured after the induction by the 645-nm (0.03 mW/cm^2^) light or in the dark for 24 hours. b) Comparison of bioluminescence intensities of mice maintained in the dark and subjected to the illumination. Data represent the mean ± SEM (n = 3 mice per group).

## Conclusion

Our work demonstrated that COMBINES-LID is efficient for creating LID systems. This method screened a generic combinatorial nanobody library using fast and cost-effective phage display and Y2H techniques to obtain high-quality, mammalian-applicable binders without need for further engineering of binding affinity and specificity, thus offering a short turnaround time (Figure S11). It relies on using protein targets with photo-induced light or dark-state specific conformational changes to select binder specificity, and theoretically can be applied to a large array of photoswitchable proteins (Table S1) to create orthogonal LID systems with diverse optical and structural properties. Applicable dimerization binders can also include those with other scaffolds such as non-immunoglobulin^[35]^ and computationally designed scaffolds.^[36]^

The LDB-3 and LDB-14-*Dr*BphP LID systems, now named ‘*nanoReD1*’ and ‘*nanoReD2*’, respectively, have simplified structures and improved *in vivo* performance, overcoming the intrinsic limitations of naturally occurring BphP LID and its derivatives. Although these systems have only been tested for light-activated gene expression, they should be useful for controlling other cellular processes, for example, the spatiotemporal activation of chimeric antigen receptor T (CAR-T) cells.^[37,38]^ Their use of the mammalian endogenous metabolite as chromophore and the compatibility of deep tissue penetration offer the unique potential to address clinical challenges such as CAR-T therapy targeting solid tumors.

## Acknowledgements

We thank Prof. N. Zheng and Dr. T. Hinds for the help on the ITC assay, Y. Wu, E. J. Quitevis and M. Moussa for the help on the nanobody screening, and T. Lin, M. Dinh and X. Kuang for the help on protein production. This work was supported by a grant from the U.S. National Institutes of Health (R35GM128918 to L.G.), a Safeway Pilot Award (to L.G.), and a startup fund of the University of Washington (to L.G.). K.W. was sup-ported by an Institute for Protein Design summer fellowship.

## Entry for the Table of Contents

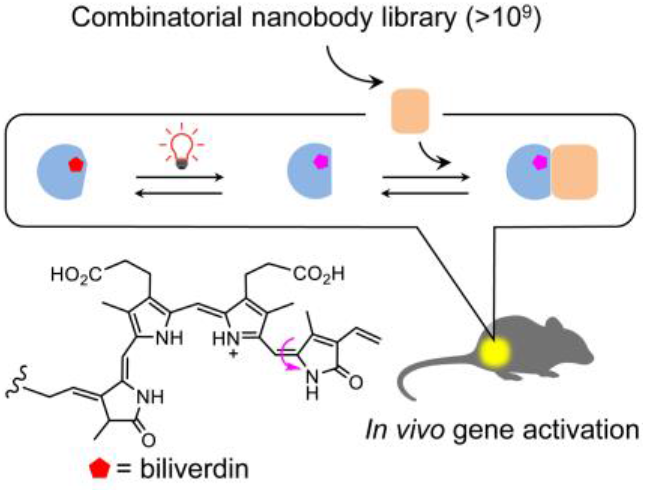

**Turning tiny antibodies into optical biosensors:**An efficient, generalizable binding protein screening method was demonstrated by creating red light-induced protein heterodimerization systems with high specificity and low dark activity, which offer deep-tissue spatiotemporal control of gene expression and other biological processes.

Institute and/or researcher Twitter usernames: @UWBiochemistry @UWproteindesign @UWgulab

