## Supporting Information for "Creating Red Light-Controlled Protein Dimerization Systems as Genetically Encoded Actuators with High Specificity"

### SUPPLEMENTARY METHODS

**Plasmid construction**

Primers and protein coding sequence (CDSs) for plasmid construction were synthesized by Integrated DNA Technologies (IDT) or amplified from other plasmids. Primers are listed in Tables S6. CDSs, noncommercial plasmid sequences, and subcloning insertion sites are listed in Table S7. The subcloning was performed using a Gibson assembly protocol recently described.[1]

**Protein expression and purification**

*Dr*BphP-Avi-His and *Dr*BphP-His were expressed in *Escherichia coli* and purified by Ni-affinity and size-exclusion chromatography. In brief, *Escherichia coli* C41(DE3) cells (Lucigen) were transformed with a *Dr*BphP-Avi-His or *Dr*BphP-His expression construct and grown in 2×YT medium at 37˚C to an $OD_{600}$ of ~0.6 before induction with 0.1% arabinose at 25˚C for overnight. Harvested cell pellets from 1-liter cultures were resuspended in 40 mL ice-cold lysis buffer (50 mM sodium phosphate, pH 8.0, 300 mM NaCl, 10 mM imidazole, 10% glycerol) for sonication. The supernatant after centrifugation at 15,000×g, 4˚C for 30 min was loaded onto a 5 mL HisTrap column (GE Healthcare) pre-equilibrated with the lysis buffer. The column was washed with a washing buffer (50 mM sodium phosphate, pH 8.0, 300 mM NaCl, 20 mM imidazole, 10% glycerol) and then His-tagged *Dr*BphP was eluted with an elution buffer (50 mM sodium phosphate, pH 8.0, 300 mM NaCl, 500 mM imidazole, 10% glycerol). Eluates were desalted with a HiPrep 26/10 column (GE Healthcare) pre-equilibrated with a storage buffer (1×PBS, 5% glycerol). Fractions were pooled and incubated with biliverdin (Frontier Scientific) with molar ratio of 1:20 at 4˚C overnight and then loaded onto a HiPrep 26/10 desalting column (GE Healthcare) pre-equilibrated with a storage buffer (1×PBS, 5% glycerol). Eluates were concentrated with Amicon Ultra-15 centrifugal filter units (30 kDa cutoff; Millipore). Concentrated proteins were loaded onto a HiLoad 16/600 Superdex 200 pg column (GE Healthcare) pre-equilibrated with a storage buffer (1×PBS, 5% glycerol). Eluted proteins were concentrated, examined by SDS-PAGE, and quantified by a Bradford assay (BioRad), then flash frozen by liquid $N_2$ and stored at -80˚C.

His-tagged or Avi-Tagged nanobodies were expressed in *Escherichia coli* strain WK6 and purified by Ni-affinity and size-exclusion chromatography as we previously reported.[2]

**Protein biotinylation**

*Dr*BphP-Avi-His was biotinylated by BirA using a BirA-500 kit (Avidity). Typically, 200 µL BiomixA (10× concentration: 0.5 M bicine buffer, pH 8.3), 200 µL BiomixB (10× concentration: 100 mM ATP, 100 mM $Mg(OAc)_2$, 500 µM d-biotin), 200 µL BIO200 (10× concentration: 500 µM d-biotin), 20 µL 1 mg/mL BirA, and *Dr*BphP-Avi-His (final concentration at ~2.4 mg/mL) were mixed with $H_2O$ to a final volume of 2 mL. The biotinylation mixture was incubated at 37˚C for 1 h and then loaded onto a HiPrep 26/10 desalting column (GE Healthcare) pre-equilibrated with a storage buffer (1×PBS, 5% glycerol). Eluted proteins were concentrated, examined by SDS-PAGE, and quantified by the Bradford assay, flash frozen by liquid $N_2$, and stored at -80˚C. LDB-3-Avi-His and LDB-14-Avi-His were biotinylated similarly as *Dr*BphP.

**Phage display selection**

The combinatorial nanobody phage library was prepared as previously described.[2] Dimerization binders were selected using 775-nm and 654-nm illuminations for the negative and positive selections, respectively. Briefly, 1.2 mL 20 µM biotinylated *Dr*BphP-Avi-His was bound to 600 µL streptavidin agarose resin (Thermo Scientific) and blocked with 1% casein and 1% BSA in 1×PBS pH 7.4 for 30 min at 4˚C in the dark. The resins were divided by a 2:1 ratio to pack the negative and positive selection columns (HR 5/5, GE Healthcare), respectively. As shown in Figure S3, both columns were connected to AKTA FPLC system and equilibrated with ~10 mL PBS buffer at 0.5 mL/min. The negative and positive selection columns were illuminated with 775 ± 14 nm light (FC-LED-780M, Prizmatix) at 0.8 mW/cm$^2$ for 10 min and 654 ± 11 nm light (FC-LED-655A, Prizmatix) at 0.3 mW/cm$^2$ for 10 min and then wrapped with aluminum foil. The light intensity was measured with an optical power meter (PM100D, Thorlabs) connected to a S130C probe (Thorlabs). In each round, phage-displayed nanobodies were loaded onto the columns equilibrated with 1× PBS at a flow rate of 0.04 mL/min. Next, the negative selection column was removed and the positive selection column washed with ~30 mL 0.05 % PBST (1×PBS with 0.05% v/v Tween 20) at a flow rate at 0.5 mL/min until the UV 280 nm baseline became stable (i.e., non-bound phages were washed out). Prior to the illumination, 2 mL flow through was collected as a "pre-elution fraction" at 0.5 mL/min immediately. The flow rate was decreased to 0 and then the positive selection column was illuminated with the 775-nm light (0.8 mW/cm$^2$) for a given time (refer to Figure S3). A 2-mL fraction was collected as a "light-elution fraction" at 0.5 mL/min immediately after the illumination. The percentage of phages specifically eluted by the light was estimated by comparing phage counts in the pre-elution and light-elution fractions. The light eluted phages were amplified and used as an input for next round biopanning.

**Y2H screening**

CDSs of the enriched nanobody library after four rounds of the biopanning were subcloned into pGADT7 to create a sub-library as preys. *Dr*BphP was inserted to pGBKT7 as the bait. Y2HGold cells were co-transformed with bait and prey plasmids, plated onto SD/-Ade/-His/-Leu/-Trp plates under the 654-nm illumination (0.03 mW/cm$^2$), and incubated at 30˚C for 4-5 days. ~2,000 well-grown clones were picked and grew in 1-mL SD/-Leu/-Trp medium in deep 96- well plates under the 654-nm illumination (0.03 mW/cm$^2$) for 24 h. 1-µL cells of each clone were replica spotted to SD/–Ade/–His/–Leu/–Trp plates and incubated under the 654-nm illumination (0.03 mW/cm$^2$) or in the dark for 2-3 days. Clones showing significantly faster growth under the illumination were picked for further analysis. Because clones picked from the plates were often contaminated with a small amount of other clones, plasmids were purified from yeast, transformed into an *E. coli* DH5α strain to select clones carrying pGADT7 on LB Agar plates with Ampicillin (100 µg/mL) and then identify those carrying correct nanobody genes by Sanger sequencing. To further confirm the gene activation specificity, sequenced preys and the bait were again co-transformed into Y2HGold cells; non-diluted and diluted (1/10 and 1/100) cells were spotted onto SD/-Ade/-His/-Leu/-Trp plates to compare colony growth under the illumination and in the dark. Sequence- and specificity-validated clones were chosen for further analyses.

**Phage ELISA**

*E. coli* electrocompetent TG1 cells were transformed with pADL-23c inserted with selected nanobody candidates. Colonies were inoculated into 250 µL media (2×TY, 2% glucose, 100 µg/mL ampicillin) in deep 96-well plates and grown at 37˚C for overnight. 10-µL cultures were inoculated into 500 µL fresh media and cells were grown to $OD_{600}$ = ~0.5 and infected by CM13 helper phage with the multiplicity of infection (MOI) of ~18. Cells were shaken at 37˚C for 45 min, added with kanamycin (50 µg/mL, the final concentration), and grown at 25˚C for overnight. Plates were centrifuged for 30 min at 3,000×g and phage-containing supernatants were transferred to fresh plates for an ELISA assay. Specifically, ELISA plates (Nunc MaxiSorp, Thermo Fisher Scientific) were coated with 100 µL 5 µg/mL streptavidin in a coating buffer (100 mM carbonate buffer, pH 8.6) at 4˚C for overnight. After washing five times with 0.05% PBST (1×PBS with 0.05% v/v

Tween 20), each well was added with 100 μL 2 μM biotinylated *Dr*BphP-Avi-His and incubated at room temperature (r.t.) for 1 h in the dark. Wells were washed five times with 0.05% PBST, blocked with 1% casein in 1× PBS, and then illuminated by the 654-nm (0.3 mW/cm$^2$) or 775-nm (0.2 mW/cm$^2$) light for 10 min. 100 μL phage supernatants were added and incubated at r.t. for 1h in dark. Wells were washed 10 times with 0.05% PBST and then illuminated with corresponding light (654 nm at 0.3 mW/cm$^2$ or 775 nm at 0.2 mW/cm$^2$) for 10 min before washing five times with 0.05% PBST. Wells were added with 100 μL HRP-M13 major coat protein antibody (RL-ph1, Santa Cruz Biotechnology; 1:10,000 dilution with 1× PBS, 1% casein) and incubated at r.t. for 1 h in the dark. A colorimetric detection was performed using a 1-Step Ultra TMB ELISA substrate solution (Thermo Fisher Scientific); $OD_{450}$ was measured with a SpectraMax Plus 384 microplate reader (Molecular Devices).

**Mammalian two-hybrid assay**

HEK293T cells (ATCC, CRL-3216) were grown in a Dulbecco's modified Eagle's medium (DMEM) supplemented with 10% fetal bovine serum (FBS; Thermo Fisher Scientific) in a humidified incubator (Forma Scientific) under 5% $CO_2$ at 37˚C. For a firefly luciferase assay, cells were grown in 24-well plates (Greiner Bio-One) to ~60% confluence and transiently co-transfected with the *Dr*BphP bait and nanobody preys, and a luciferase reporter plasmid (Addgene, #64125) in a 1:1:1 ratio (0.25 μg each into a ~500 μL medium). After the transfection, culture medium was changed in 6 h and then cells were kept in the darkness for another 18 h prior to the transcription activation. The activation was performed by continuously illuminating cells with the 654-nm light at 0.2 mW/cm$^2$ for 24 h; cells were kept in the dark as the control. The time-course luciferase assay was performed as described above with different transcription induction times.

Luciferase levels were measured with a firefly luciferase glow assay kit (Pierce) following the manual. Briefly, after the transcription activation, cells were washed with 1× PBS, added with 150 µl of 1× cell lysis buffer, and incubated at 4˚C for 30 min. 20 µl cell lysate from each well was transferred into a black 96-well plate (CELLSTAR, Greiner Bio-One, Cat # 655079) and mixed with 50 µl of a Working Solution. Bioluminescence signals were measured with a SpectraMax i3 plate reader (Molecular Devices) after incubation in the dark at r.t. for 10 min.

The luciferase assay of *Rp*BphP1-based systems was performed under the same condition, except that the transcription activation was performed with the 775-nm (0.2 mW/cm$^2$) illumination, because different from *Dr*BphP, *Rp*BphP1 is converted to the light form by NIR illumination.

#### Analysis of nanobody stability in mammalian cells

~2×10$^5$ HEK293T cells were seeded in 6-well plates (Thermo Fisher Scientific, catalog # 140675) in DMEM supplemented with 10% FBS, and incubated under 5% $CO_2$ at 37˚C for overnight. Cells in a 1.5-mL medium were transiently transfected with plasmids (2.5 μg each) encoding nanobody–SNAP-tag fusions using lipofectamine 2000 (Thermo Fisher Scientific). After 36-h incubation, the medium was removed and cells were washed with 1× PBS twice, dissociated from the plate by digestion with a 1× Trypsin-EDTA Solution (Thermo Fisher Scientific, catalog # R001100), and collected in 15-mL conical tubes. Cells were washed with 1 mL 1× PBS and re-suspended in 250 μL ice cold 1× PBS for sonication. After centrifugation at 20,000g for 10 min, ~50 μL supernatants were incubated with 1 μM (final concentration) SNAP-Surface 649 (New England Biolabs, catalog # S9159S) for 1 h at r.t. to label SNAP-tagged proteins. Labelled samples were boiled for 10 min at 95˚C in an SDS sample loading buffer before loaded onto an SDS-PAGE gel. The gel was scanned by an Odyssey CLx imaging system (Li-cor Biosciences).

#### Analytical SEC

Interactions of *Dr*BphP with LDB-3 and LDB-14 were analyzed by analytical SEC. Samples were loaded onto a Superdex 200 Increase 10/300 GL column (GE Healthcare) pre-equilibrated with 1× PBS and eluted at 0.75 mL/min at 4˚C. The column was calibrated with molecular weight standards (Bio-Rad, catalog # 1511901). Light-sensitive samples were prepared in a dark room and the column and sample syringes were all covered by aluminum foil to avoid light exposure.

To detect the complex formation, *Dr*BphP-His was photoconverted to the dark and light forms by the 775-nm (0.8 mW/cm$^2$, 10 min) and 654-nm (0.2 mW/cm$^2$, 5 min) illumination, respectively. ~6 μM (final concentration) *Dr*BphP-His was added with ~5 μM (final concentration) LDB-3-His or LDB-14-His and incubated at r.t. for 30 min in the dark before loading a 500 μL mixture onto the column. 500-μL fractions with an elution volume between 8 and 16 mL were collected and proteins in each fraction were precipitated by trichloroacetic acid (TCA) for SDS-PAGE analysis. Briefly, 55 μL 100% TCA was mixed with each fraction and incubated at -20˚C for 30 min. After centrifugation at 20,000×g, 4˚C for 15 min, supernatants were removed, and pellets were washed with 600 μL ice-cold acetone three times and then dried in air. Pellets were resuspended and boiled in the SDS loading buffer and analyzed by SDS-PAGE.

#### Isothermal titration calorimetry

Binding affinities and thermodynamics of LDB-3 and LDB-14 to *Dr*BphP were measured by a MicroCal PEAQ-ITC device (Malvern) at 25˚C. Specifically, ~ 210 μL of 10 or 5 μM *Dr*BphP-His was loaded to a sample cell and then illuminated by the 654-nm (0.2 mW/cm$^2$, 5 min) or 775-nm (0.8 mW/cm$^2$, 15 min) light. ~38 μL 80 μM LDB-3-His and 50 μM LDB-14-His were titrated into 10 and 5 μM *Dr*BphP-His in the cell, respectively, by 19 injections (2 μL each) from a syringe. Background heat transfer caused by the nanobody dilution was measured by conducting a titration of LDB-3-His (80 μM) or LDB-14-His (50 μM) into a 1× PBS buffer alone. Titration of 1× PBS buffer into *Dr*BphP-His (10 μM) was also conducted as the control.

Raw ITC data were analyzed by NITPIC version 1.2.7.[3] To find a suitable range for each injection, cut-off differentials for the injection end was changed to 0.1. The fitting equation for a one-site model is $y = \frac{L}{1+e^{-k(x-x_0)}} + b$, where $y$ represents the heat of injection, $x$ represents the molar ratio, and $b, k,\ L,\ x_0$ are related parameters. The integrated data of LDB-3 and LDB-14 titrated to the light form were fitted with the above equation by using a "curve_fit" function in the Python-SciPy package, which generated $K_D^{app}$ and other thermodynamic parameters in Table S4.

**Bio-layer interferometry**

LDB-3 and LDB-14 binding kinetics were analyzed using an Octet RED96 system (ForteBio) and Streptavidin (SA) biosensors. Briefly, 20 μg/mL biotinylated LDB-3-Avi-His or LDB-14-Avi-His was immobilized on SA biosensors in 1× PBS buffer (pH 7.4). A duplicate set of sensors was incubated in the buffer without any protein to measure background binding. All sensors were blocked with a buffer (1× PBS, pH 7.4, 0.05% Tween-20, 0.2% BSA, and 10 ng/mL biocytin) before the binding assay. Serial dilutions of *Dr*BphP-His in an assay buffer (1× PBS, pH 7.4, 0.05% Tween-20, and 0.2% BSA) were illuminated with the 654-nm (0.3 mW cm$^{-2}$, 5 min) or 775-nm (0.2 mW cm$^{-2}$, 10 min) light before binding to the nanobodies. The assay was performed in black 96-well plates with a total working volume of 0.2 mL per well at r.t. Raw data were analyzed by an Octet data analysis software V9.0 (ForteBio) using a double-reference-subtraction protocol to subtract signals related to nonspecific binding, background, and signal drift caused by sensor variability.

Apparent dissociation constants ($K_D^{app}$s) were calculated by the steady-state analysis and the fitting with a global 1:1 model. The fitting of apparent dissociation rate constant ($k_{off}^{app}$) was found to be more reliable (or less *Dr*BphP-His concentration dependent) than the fitting of apparent binding rate constant ($k_{on}^{app}$), so only $k_{off}^{app}$ was calculated by fitting with the equation, $C = C_0 + A(1 - e^{-k_{off}t})$, where $C$ represents the level of binding, $C_0$ the binding at the start of dissociation, $A$ an asymptote, and t time. $k_{off}^{app}$ for each binding was calculated using the "curve_fit" function in the Python-SciPy package. After obtaining $K_D^{app}$ and $k_{off}^{app}$, $k_{on}^{app}$ was calculated by $k_{on} = \frac{k_{off}}{K_D}$. Of note, compared with the fitting result, the dissociation curves were slightly tailed (Figure S8), likely due to the contribution from *Dr*BphP dimer dissociation.

##### *Dr*BphP photoconversion analysis

The *Dr*BphP thermal relaxation efficiency was analyzed by absorption spectroscopy. Absorption spectra (500-900 nm) of *Dr*BphP samples were obtained using a SpectraMax Plus 384 microplate reader (Molecular Devices). *Dr*BphP-His was added in a quartz micro cuvette (Yixing Purshee Optical Elements) and then converted to the light or dark form by the 654-nm (0.5 mW/cm$^2$, 2 min) or 775-nm (0.3 mW/cm$^2$, 10 min) illumination before collecting spectra. To monitor the real-time thermal relaxation to the dark form, ~400 μL 5 μM (final concentration) *Dr*BphP-His samples added with or without 5 μM (final concentration) LDB-3-His or LDB-14-His in the cuvette were first converted to the light form by the 654-nm (0.5 mW/cm$^2$) illumination for 2 min and then immediately relaxed by the 775-nm (0.3 or 0.05 mW/cm$^2$) illumination with different exposure times before collecting spectra. The ratio of $A_{750}/A_{700}$ was normalized to the range (0-1) to monitor the relaxation process.

##### GFP imaging

HEK293T cells were seeded in 10 cm Nunclon Delta Surface culture dishes (Thermo Scientific) in DMEM supplemented with 10% FBS in the humidified incubator under 5% $CO_2$ at 37˚C. They were co-transfected with 10 μg a pGreenFire1-Gal4 lentivector (System Biosciences, catalog # TR017PA-1) and lentivirus-packing plasmids (5μg each) including PMDL, REV and VSVG by a calcium phosphate transfection method. The medium was changed in 6 h after the transfection and the virus was harvested after incubation for another 72 h. To separate the virus from the medium, the medium was centrifuged at 500×g for 5 min and the supernatant was passing through a Millex-HV filter (0.45 μm, Merck Millipore). 2.5 out of 10 mL filtered virus was used to infect HEK293T cells cultured in another 10 cm dish under 50% confluence, with 10 μg/mL polybrene (Merck Millipore), for 24 h.

Lentivirus-transduced HEK293T cells were seeded in 35 mm glass bottom microwell dishes coated with poly-D-lysine (MatTek, catalog # P35GC-0-10-C) at a density of $1\times10^5$ cells per dish. On the second day, cells were transiently co-transfected with the GAL4-BD–*Dr*BphP and nanobody-p65 plasmids (1.25 μg each) using lipofectamine 2000 (Thermo Fisher Scientific) and incubated for overnight. For each nanobody candidate, two dishes were needed for the illumination and the dark control; after the transfection, dishes were immediately covered by aluminum foil to avoid light exposure. On the third day, cells were under the 654-nm (0.2 $mW/cm^2$) illumination or maintained in the dark for another 48 h. Prior to fluorescence imaging, cells were fixed by 4% paraformaldehyde for 10 min and washed with 1× PBS.

GFP images were acquired using a Nikon Ti-E automated inverted microscope equipped with a Perfect Focus System, a Nikon 20×/0.75-NA Plan Apo Lambda objective, a linear encoded motorized stage (Nikon Ti-S-ER), and an Andor iXon Ultra 888 EMCCD camera (16-bit dynamic range, 1,024×1,024 array with 13-μm pixels). Cells were illuminated by a SPECTRA X LED illuminator (Lumencor) coupled with an excitation filter (448 ± 19 nm) and a filter cube mounted with a dichroic mirror (506 nm) and an emission filter (510 ± 20 nm) (Chroma). Culture dishes were scanned under the GFP and a brightfield channels. Acquired GFP images (dark and light condition) were analyzed by MATLAB for quantifying the fluorescence intensity. Specifically, fluorescence signals in all pixels were subtracted by an average background value (i.e., the median of the pixel intensity distribution in each field-of-view (FOV)) and integrated for each FOV. For each condition, 78 FOVs were sampled for statistical analysis.

**Transcription activation in mice**

6- to 8-week-old male BALB/c nude mice of ~20 g body weight (the Jackson Laboratory) were used for the *in vivo* gene activation controlled by the *Dr*BphP–LDB-3 LID system. Mice were subcutaneously injected with HEK293T cells co-transfected with the plasmids encoding LDB-3-p65, GAL4-DB–*Dr*BphP, and the luciferase reporter. Specifically, each mouse was injected with $\sim5\times10^6$ cells supplemented with 100 μL DMEM medium 6 h after the transfection, and kept in a conventional cage in the darkness for 24 h. In another 24 h, mice in a light treatment group (n = 3) were placed in a transparent cage illuminated by a 654 nm LED array at 0.03 $mW/cm^2$ and those in the control group (n = 3) remained in the dark. Mice were subjected to bioluminescence imaging using an IVIS Spectrum instrument (PerkinElmer) in luminescence mode with an open emission filter. Throughout the imaging process, animals were maintained under anesthesia with 1.5%

vaporized isoflurane. Prior to imaging, 200 μL of 15 mg/mL D-Luciferin sodium salt solution (US Everbright) was intravenously injected through a tail vein. Data were analyzed using a Living Image 3.0 software (Perkin Elmer). Animal data were collected in Third Institute of Oceanography following a protocol approved by the University of Washington Institutional Animal Care and Use Committee (IACUC).

### SUPPLEMENTARY FIGURES

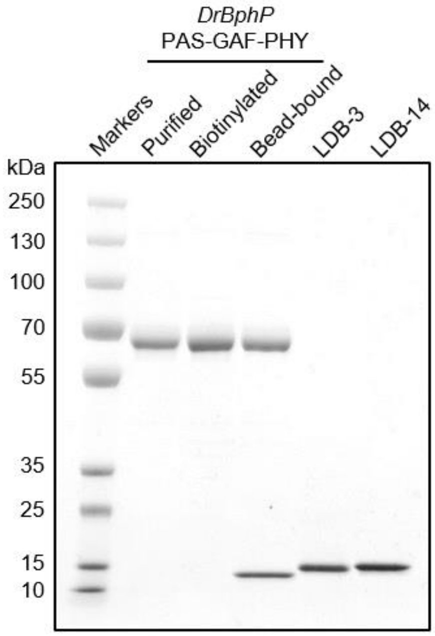

**Figure S1.** SDS-PAGE analysis of purified *Dr*BphP and nanobodies. Proteins were purified by nickel affinity and SEC chromatography. To examine *in vitro* biotinylation efficiency by BirA, the biotinylated protein was bound to streptavidin beads (Dynabeads M-280 Streptavidin, Thermo Fisher Scientific) and the bound protein (lane 4) was compared with the input protein (lane 3). The lower band in the lane 4 was streptavidin released from the beads when boiling the sample in an SDS loading buffer.

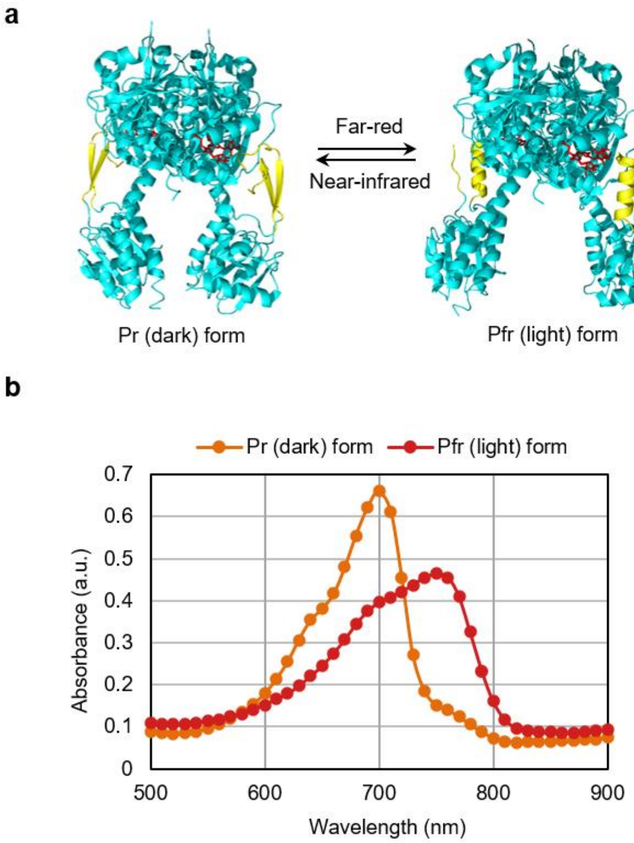

**Figure S2.** Structures and spectra of the dark and light forms of *Dr*BphP. (a) Structures of *Dr*BphP dark and light forms previously reported[4] showing the biliverdin chromophore (red sticks) bound to a tri-domain photosensory module (cyan cartoon) and conformational changes of a tongue motif (yellow) interacting with the biliverdin binding pocket. (b) Absorption spectra of the dark and light states of the biotinylated *Dr*BphP after the 775-nm (0.3 mW/cm$^2$, 10 min) and 654-nm (0.5 mW/cm$^2$, 2 min) illuminations, respectively.

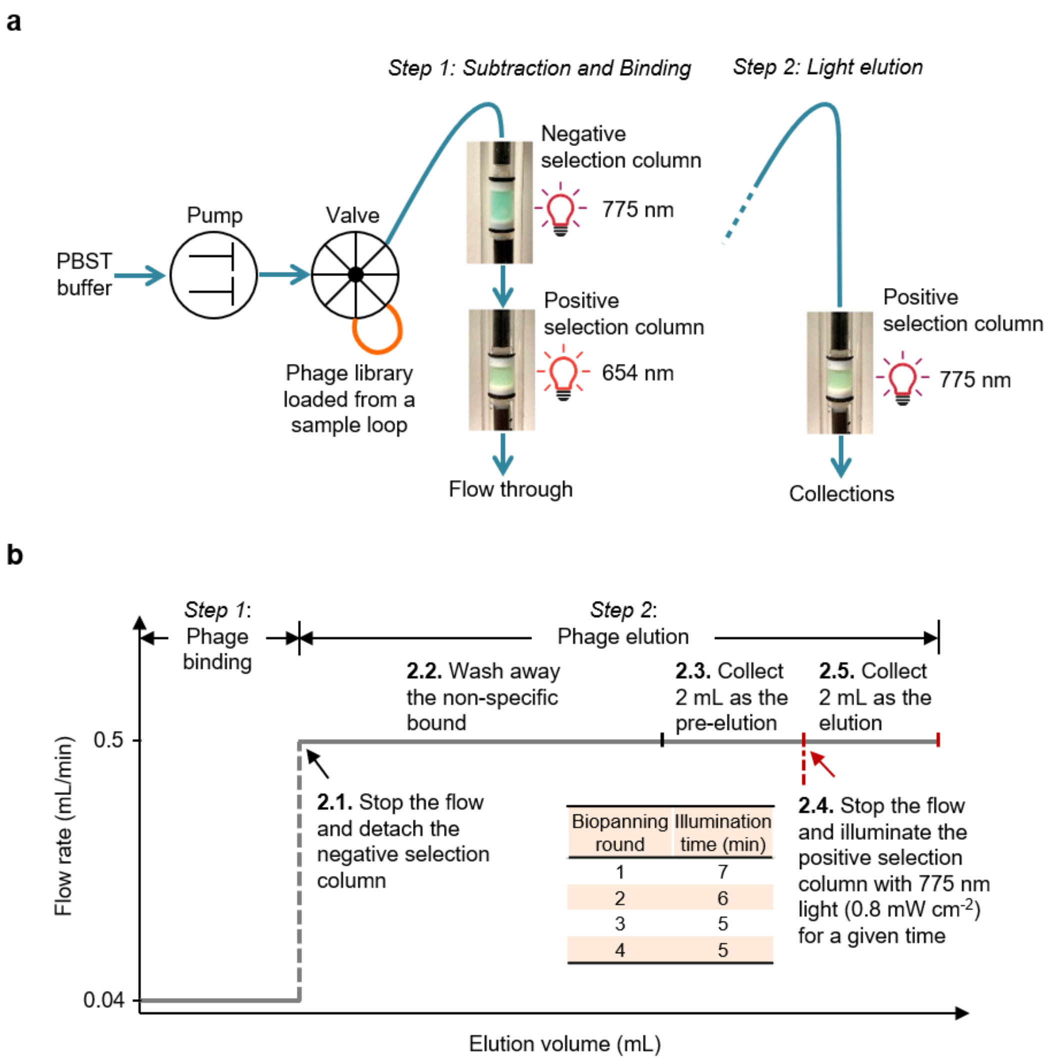

**Figure S3**. Column chromatography-based phage display selection. (a) Two-step biopanning FPLC setup. (b) Flow rate and illumination time setup. In the Step 1, 2 mL phage-displayed nanobodies were loaded to two connected transparent glass columns (HR 5/5, GE Healthcare) packed with 0.4 and 0.2 mL streptavidin agarose resin (Pierce). Before divided into the two columns, the resin was incubated with 1.2 mL 20 µM biotinylated *Dr*BphP in the dark for 30 min. Next, *Dr*BphP in the first (negative selection) and second (positive selection) columns were converted to the dark and light forms by the 775-nm (0.3 mW/cm$^2$, 10 min) and 654-nm (0.5 mW/cm$^2$, 2 min) illumination, respectively. After the phage injection, the flow rate was set to be 0.04 mL/min and then decreased to 0 when the UV 280 nm baseline was stable (i.e., non-bound phages were washed out). In the Step 2, the first column was removed, and phages were eluted from the second column by the 775-nm (0.8 mW/cm$^2$) illumination for a given time. A pre-elution fraction was collected as a control for the phage count comparison with a light elution fraction to estimate the ratio of phages specifically eluted by the light to those non-specifically eluted (refer to Table S2).

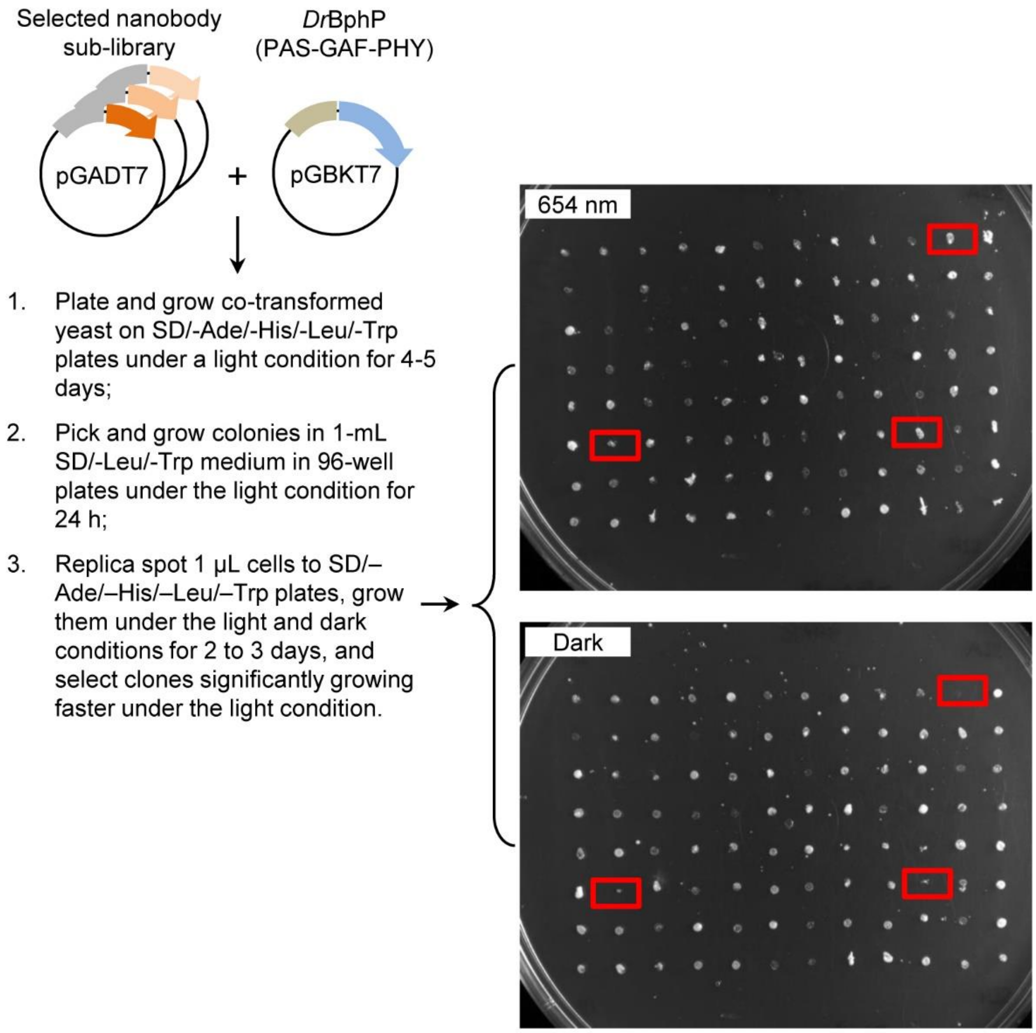

**Figure S4**. Yeast two-hybrid screening. Phage display-enriched nanobodies (orange), as preys, were subcloned to pGADT7 encoding a GAL4 AD domain (grey). *Dr*BphP (blue), as a bait, was inserted to pGBKT7 encoding a GAL4 DNA-binding domain (green). The right panel shows a representative result of two replica spotted plates incubated in the dark or under the 654-nm illumination.

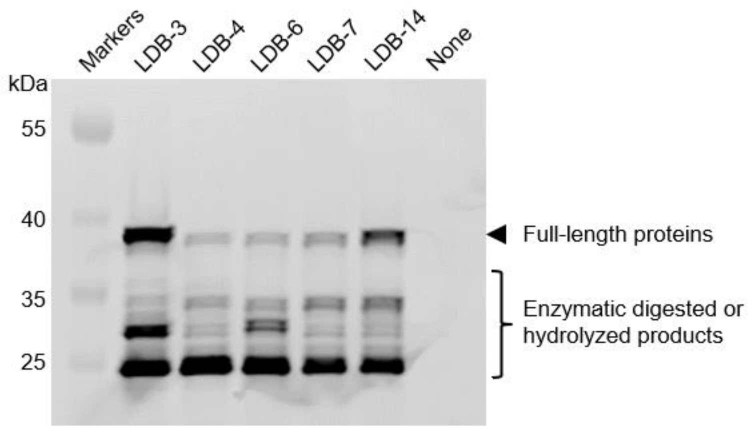

**Figure S5**. In-gel detection of fluorescently labelled nanobodies expressed in HEK293T cells. Cells were transiently transfected with plasmids encoding SNAP-tagged nanobody fusions. Proteins in supernatants of sonication-lysed cells were specifically labeled with SNAP-Surface 649 and analyzed by SDS-PAGE and fluorescence imaging with an Odyssey CLx imaging system.

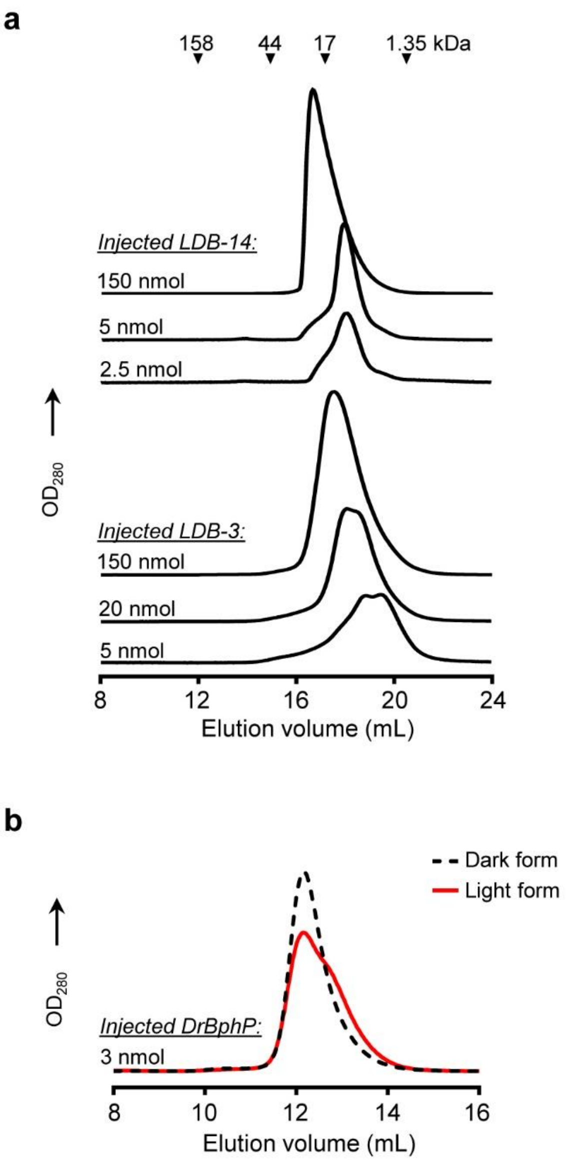

**Figure S6.** Analytical SEC analyses of (a) nanobodies at different concentrations and (b) *Dr*BphP in the light and dark forms. Proteins were loaded to a Superdex 200 Increase 10/300 GL column pre-equilibrated with 1× PBS and eluted at a flow rate of 0.75 mL/min at 4˚C.

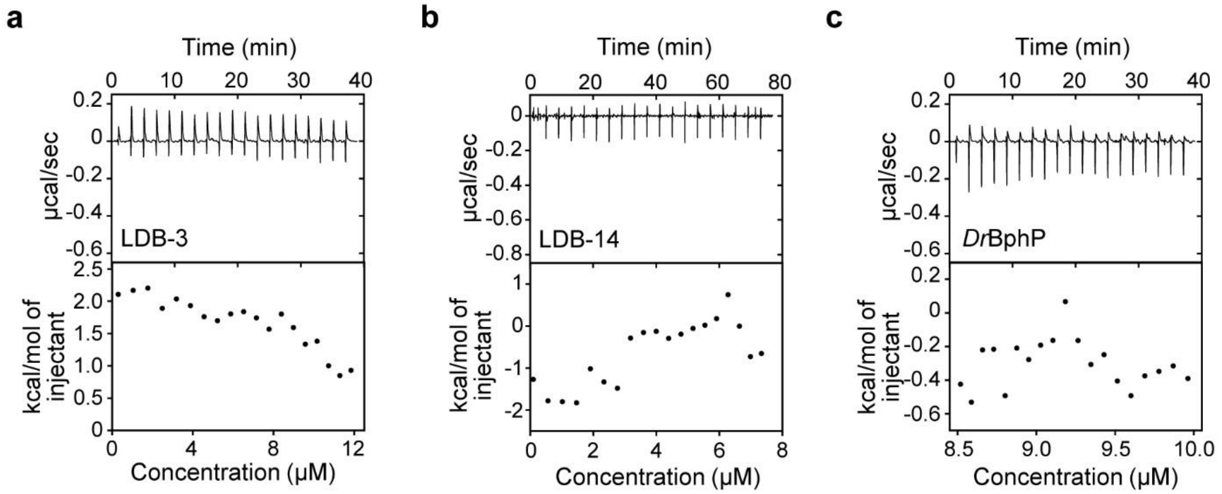

**Figure S7**. ITC analysis of the titration of (a) 80 μM LDB-3 or (b) 50 μM LDB-14 into 1× PBS buffer, and (c) the titration of 1× PBS buffer into 10 μM *Dr*BphP. The raw data (top) and the integration of heats (bottom) for each titration are shown.

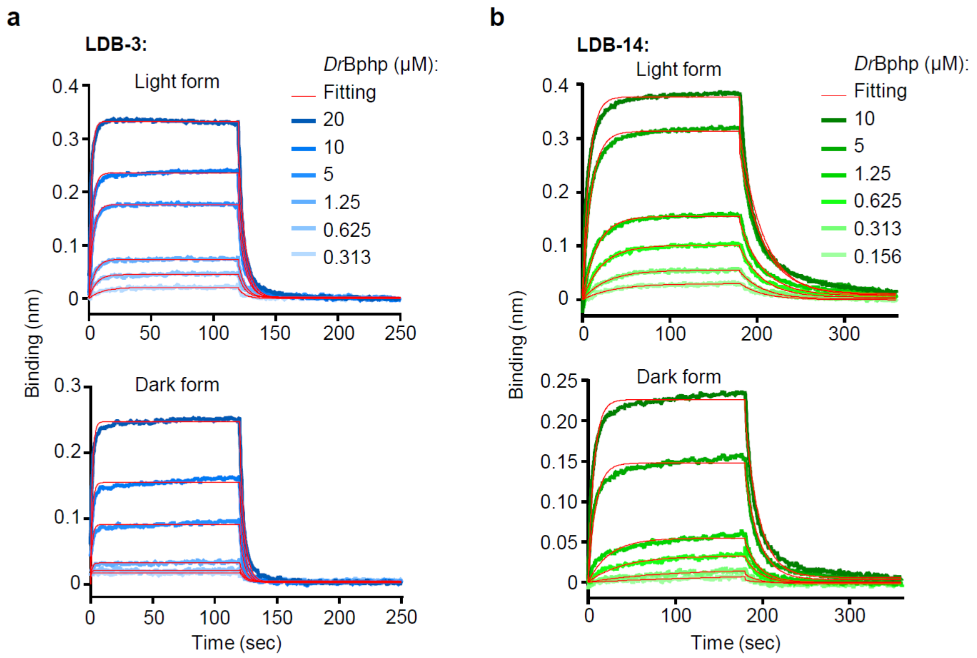

**Figure S8.** BLI analysis of LDB-3 and LDB-14 binding kinetics. BLI sensorgrams show *Dr*BphP binding to LDB-3 (a) and LDB-14 (b). Nanobodies were immobilized on Streptavidin biosensors and interacted with *Dr*BphP after the 654-nm (light form) or 775-nm (dark form) illumination. Data were fitted using a global 1:1 model.

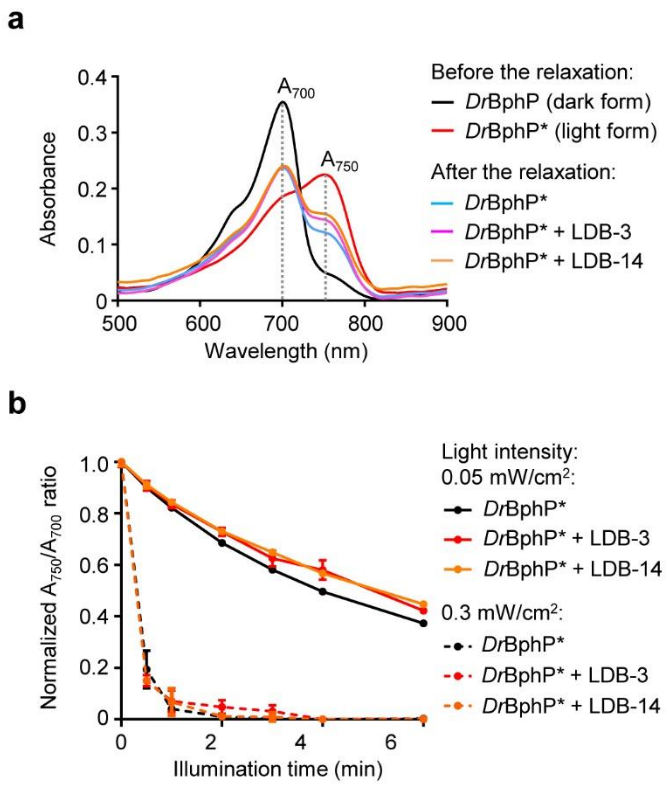

**Figure S9.** Inhibition of *Dr*BphP relaxation to the dark form by the nanobody binding. (a) Representative absorption spectra of the photoconverted light and dark forms and after the thermal relaxation by 775-nm illumination with or without LDB-3 or LDB-14 binding. (b) Time-course analysis of thermal relaxation rates of unbound and nanobody-bound light-form *Dr*BphP by the 775-nm illumination. 400 µl 5 µM (final concentration) light-form *Dr*BphP (after the 654-nm illumination at 0.5 mW/cm$^2$ for 2 min) was incubated with 5 µM (final concentration) LDB-3 or LDB-14 for 10 min before the thermal relaxation.

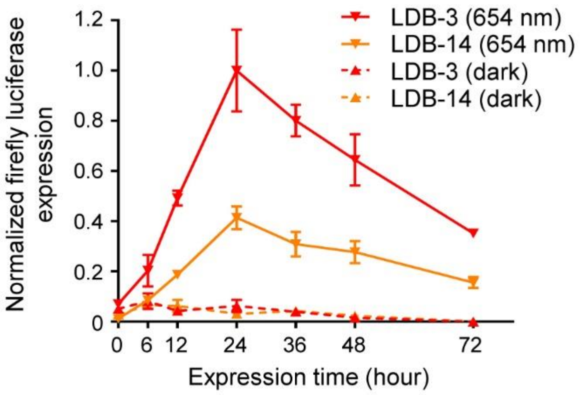

**Figure S10.** Time-course analysis of red light-induced luciferase expression. HEK293T cells were co-transfected with the bait, prey, and GAL4UAS-luciferase reporter plasmids (~0.25 μg each) in a 0.5 mL culture. Transfected cells were incubated under the 654-nm (0.2 mW/cm$^2$) illumination or in the dark. Data represent mean values of 3 measurements; error bars, standard deviation.

1. Production of a biotinylated conformation switcher (*2-3 weeks*)

↓

2. Phage display selection of a combinatorial binder library to obtain a sub-library of enriched dimerization binders (typically 2 to 4 rounds, *1-2 weeks*)

↓

3. Yeast two-hybrid screening of the sub-library to obtain dimerization binder hits (*3-4 weeks*)

↓

4. Expression and purification of dimerization binder hits (*1-2 week*)

↓

5. *In vitro* binding specificity validation by ELISA or other methods (*2 days to 1 week*)

↓

6. Functional validation of biosensor candidates by a mammalian two-hybrid assay (*~1 week*)

↓

Test LID biosensors for applications

**Figure S11**. Flowchart and timeline of COMBINES-LID.

### SUPPLEMENTARY TABLES

**Table S1.** Summary of photoswitchable proteins or domains that can potentially be used as conformational switchers in LID systems.

| Photo-switchable proteins (or domains) | Example(s) | Chromophore(s) | Excitation λ (nm) | Reversion λ (nm) | Oligomeric state | | Natural light induced binder(s) | Reference (s) |
|---|---|---|---|---|---|---|---|---|
| | | | | | Dark form | Light form | | |
| **UV receptors** | UVR8 | Trp | ~300 | Dark | Dimer | Monomer | COP1 | 5-7 |
| **Cyanobacteriochromes** | CcaS | PCB | ~535 | ~672/dark | Monomer | Monomer | CcaR | 8-9 |
| | cPAC | PCB | ~410 | ~520/dark | Dimer | Dimer | Unknown | 10 |
| | UirS | PCB | ~400 | ~530/dark | Monomer | Monomer | UirR | 11-12 |
| **Sensors of blue-light using FAD (BLUF) domains** | PixD | FAD or FMN | ~450 | Dark | Decamer | Dimer | PixE | 13-15 |
| | bPAC | FAD or FMN | ~450 | Dark | Dimer | Dimer | Unknown | 16-18 |
| **LOV domains** | AsLOV2 | FMN | ~450 | Dark | Monomer | Monomer | Unknown | 19-21 |
| | YtvA | FAD, FMN, or riboflavin | ~450 | Dark | Dimer | Dimer | Unknown | 22-24 |
| | VVD | FAD or FMN | ~450 | Dark | Monomer | Dimer | VVD | 25-27 |
| | FKF1 | FMN | ~450 | Dark | Monomer | Dimer | GI | 28-30 |
| | EL222 | FMN | ~450 | Dark | Monomer | Dimer | Unknown | 31-32 |
| **Cryptochromes** | CRY2 | FAD | ~450 | Dark | Monomer | Monomer | CIB1 | 33-34 |
| **Fluorescent protein domains** | Dronpa145K/N | p-HBI | ~400 | ~500/dark | Monomer | Dimer | Unknown | 35-36 |
| | PYP | p-coumaric acid | ~450 | Dark | Monomer | Monomer | Unknown | 37-38 |
| **Opsins** | BeCyclOp | Retinal | ~530 | Dark | Dimer | Dimer | Unknown | 39 |
| **Cobalamin binding domains (CBDs)** | TtCBD | AdoCbl, MetCbl, or CNCbl | ~545 | Dark | Tetramer | Monomer | Unknown | 40 |
| | MxCBD | AdoCbl, MetCbl, or CNCbl | ~545 | Dark | Tetramer | Monomer | Unknown | 40 |
| **Phytochromes** | *Rp*BphP1 | BV | ~740 | ~636/dark | Dimer | Dimer | PpsR2 | 41-42 |
| | *Dr*BphP | BV | ~655 | ~780/dark | Dimer | Dimer | Unknown | 4, 43-44 |
| | Cph1 | PCB or PΦB | ~657 | ~731/dark | Dimer | Dimer | Unknown | 45-47 |
| | PhyB | PCB | ~660 | ~740/dark | Monomer | Monomer | PIF3/PIF6 | 48-50 |

**Table S2.** Enrichment of phage titers following each round of biopanning for the dimerization binder selection.

| Round | Input count | Pre-elution count* | 775 nm light elution count** |
|---|---|---|---|
| 1 | ~$1×10^{14}$ | ~$5.4×10^{6}$ | ~$2.0×10^{6}$ |
| 2 | ~$1×10^{13}$ | ~$1.2×10^{5}$ | ~$2.3×10^{5}$ |
| 3 | ~$1×10^{13}$ | ~$1.8×10^{5}$ | ~$7.3×10^{5}$ |
| 4 | ~$1×10^{13}$ | ~$1.5×10^{7}$ | ~$1.4×10^{8}$ |

**Note:** After phage binding, the positive selection column was washed with ~30 mL 0.05% PBST buffer. 2 mL pre-elution fraction (*****) was collected at 0.5 mL/min immediately before the 775 nm-illumination at 0 mL/min. 2 mL elution fraction (******) was collected at 0.5 mL/min immediately after the illumination. Phage titers of the collected fractions were measured to determine the enrichment of clones specifically eluted by the light for each selection round.

**Table S3.** CDR sequences of light-induced dimerization binders (LDBs) characterized in the work.

| Nanobody | CDR1 | CDR2 | CDR3 |
|---|---|---|---|
| **LDB-3** | FTWDHYI | ENGDAWN | IGFDVPSGRSWQGSHFWM |
| **LDB-4** | DTSYLYS | WWWNLTQ | WSIYFPPGNDYNGYH |
| **LDB-6** | FFSNWSD | FWADGTE | WYGPVNGFYMFD |
| **LDB-7** | STSDFES | SWFTNPP | HRSIWYHPT |
| **LDB-14** | TTSRWES | WQNNSVP | AQHNFLGHR |

**Table S4.** ITC-derived thermodynamic parameters for LDB-3 and LDB-14 binding to the *Dr*BphP light form.

| | **n (stoichiometry)** | **$K_D^{app}$ (μM)** | **ΔH (kJ/mol)** | **ΔG (kJ/mol)** | **-TΔS (kJ/mol)** |
|---|---|---|---|---|---|
| **LDB-3** | 0.605 | 1.01 | -37.0 | -34.2 | 2.78 |
| **LDB-14** | 0.556 | 0.47 | -112.8 | -36.1 | 76.7 |

**Table S5.** Kinetic parameters of selected dimerization binders binding to the light and dark forms.

| | LDB-3 | | | |
|---|---|---|---|---|
| | **Molar ratio** | **$K_D^{app}$ ($10^{-6}$M)** | **$k_{on}^{app}$ ($10^4$ $M^{-1}$ $s^{-1}$)** | **$k_{off}^{app}$ ($10^{-2}$ $s^{-1}$)** |
| **Light form** | 1:1 | 7.7 | 2.4 | 18.5 |
| **Dark form*** | 1:1 | 25 | 1.06 | 26.5 |
| | **LDB-14** | | | |
| | **Molar ratio** | **$K_D^{app}$ ($10^{-6}$M)** | **$k_{on}^{app}$ ($10^4$ $M^{-1}$ $s^{-1}$)** | **$k_{off}^{app}$ ($10^{-2}$ $s^{-1}$)** |
| **Light form** | 1:1 | 2.4 | 1.56 | 3.74 |
| **Dark form*** | 1:1 | 10 | 0.607 | 6.07 |

**Note:** *The dark form binding data are not reliable because the white light conducted to BLI biosensors might partially convert *Dr*BphP to the light form.

**Table S6.** Synthetic oligos used for plasmid construction in this study.

| | Vector backbone | Name | Sequences (5' to 3') | Note |
|---|---|---|---|---|
| *Dr*BphP-Avi-His | pBAD | *Dr*BphP-Avi-His F | CTTTAAGAAGGAGATAT*GGATCC*ATGAGTCGTGACCCTTTGCCAT | *Bam*HI or *Eco*RI sites are underlined. |
| | | *Dr*BphP-Avi-His R | TGGTGATGGTGATGATG*GAATTC*TTAGTGATGGTGGTGATGATG | |
| *Dr*BphP-His | pBAD | *Dr*BphP-His F | CTTTAAGAAGGAGATAT*GGATCC*ATGAGTCGTGACCCTTTGCCA | *Bam*HI or *Eco*RI sites are underlined. |
| | | *Dr*BphP-His R | TGGTGATGGTGATGATG*GAATTC*TAATGCGCCAGTAAGAGTGTC | |
| Nanobody-His | pADL-23c | Nanobody-His F | GGATTGTTATTACTCGCG*GCCCAGCCGGC*CATGGCAGAAGTTCAGCTGCAGGCAAGCGG | *Bgl*I sites are underlined. |
| | | Nanobody-His R | TGATGGTGGTGATGGTGTTG*GCCTCCCGGGC*TGCTGCTAACGGTAACCTGGGTGC | |
| Nanobody-Avi-His | pADL-23c | Nanobody-Avi-His F | TATTACTCGCG*GCCCAGCCGGC*CATGGCAGAAGTTCAGCTGCAGGCAAGC | *Bgl*I sites are underlined. |
| | | Nanobody-Avi-His R | GGTGGATAAGCTTTG*GCCTCCCGGGC*TGCTGCTAACGGTAACCTGGGTGC | |
| *Dr*BphP-Yeast | pGBKT7 | *Dr*BphP-Yeast F | GAGGAGGACCTG*CATATG*GGAGGCGGTTCCGGTGGCGG | *Nde*I or *Bam*HI sites are underlined. |
| | | *Dr*BphP-Yeast R | CTGCAGGTCGAC*GGATCC*CTAGCTGCTAACGGTAACCTGGG | |
| Nanobody-Yeast | pGADT7 | Nanobody-Yeast F | GATTACGCT*CATATG*GGAGGCGGTTCCGGTGGCGGTTCTGAAGTTCAGCTGCAGGCAAGC | *Nde*I or *Bam*HI sites are underlined. |
| | | Nanobody-Yeast R | CTCGAGCTCGAT*GGATCC*CTAGCTGCTAACGGTAACCTGGGT | |
| *Dr*BphP-Mammalian | pBobi | *Dr*BphP-Mammalian F | GTTGCCACCATG*GGATCC*ATGAAGCTACTGTCTTCTATC | *Bam*HI or *Xho*I sites are underlined. |
| | | *Dr*BphP-Mammalian R | GGAACCACCACC*CTCGAG*TAATGCGCCAGTAAGAGTGTC | |
| Nanobody-Mammalian | pBobi | 3*NLS F | GTTGCCACCATG*GGATCC*CCCAAGAAGAAGCGCAAGGT | *Bam*HI or *Xho*I sites are underlined. |
| | | 3*NLS R | TGCCTGCAGCTGAACTTCTCCGCTGCCACCAGACCCTC | |
| | | Nanobody F | GAAGTTCAGCTGCAGGCAAGC | |
| | | Nanobody R | ACTGCCACCGCCGCCGCTGCTGCTAACGGTAACCTGGGT | |
| | | p65 F | AGCGGCGGCGGTGGCAGTCAGTACCTGCCAGATACAGAC | |
| | | p65 R | GGAACCACCACC*CTCGAG*GGAGCTGATCTGACTCAGCAG | |
| LDB-14-Mammalian | pcDNA3 | 3*NLS F | AAGCTGGCTAGTT*AAGCTT*ATGCCCAAGAAGAAGCGCAAGGTG | *Hin*dIII or *Xho*I sites are underlined. |
| | | 3*NLS R | TCCGCTGCCACCAGACCCTC | |
| | | Nanobody F | GTTGAAGCATCTGGATCCGGAGGCGGTTCCGGTGGCGG | |
| | | Nanobody R | GCCACTTCCTCCGGTACCGCTGCTAACGGTAACCTGGGT | |

| | | | | |
|---|---|---|---|---|
| | | p65 F | GGGTCTGGTGGCAGCGGACAGTACCTGCCAGATACAGACGAT | |
| | | p65 R1 (first round PCR) | TCCACTGCCGCCAGAGCTGCCACTTCCTCCGGAGCTGATCTGACTCAGCAG | |
| | | p65 R2 (second round PCR) | CGGGCCCTCTAGA*CTCGAG*CTACTGAATTCTCCACTGCCGCCAGAGCTGC | |
| *Rp*BphP1-Mammalian | pcDNA3 | GAL4 BD F | ACCCAAGCTGGCTAGTT*AAGCTT*ATGAAGCTACTGTCTTCTATCG | *Hin*dIII or *Xho*I sites are underlined. |
| | | GAL4 BD R | CATATGCAGGTCCTCCTCTGA | |
| | | *Rp*BphP1-F | GAGGAGGACCTGCATATGGTGGCAGGTCATGCCTCTGGC | |
| | | *Rp*BphP1-R | GGGCCCTCTAGA*CTCGAG*CTACTTCTTGTTCTGCGAGCCATT | |
| PpsR2-Mammalian | pcDNA3 | PpsR2 F | GTTGAAGCATCT*GGATCC*GTGGCGTCAAAGTCCGTTCAT | *Bam*HI or *Kpn*I sites are underlined. |
| | | PpsR2 R | GCCACTTCCTCC*GGTACC*ATCCTCTGCGTCGTCTGAG | |
| Q-PAS1-Mammalian | pcDNA3 | Q-PAS1 F | GTTGAAGCATCT*GGATCC*GGCAAGAACATGCAGGCGGT | *Bam*HI or *Kpn*I sites are underlined. |
| | | Q-PAS1 R | GCCACTTCCTCC*GGTACC*GTCGTCGATCGCGGGAGTCG | |
| Nanobody-SNAP | pBobi | Nanobody F | ACTGAGCTCCTTAAGGTTGCCACCATG*GGATCC*GAAGTTCAGCTGCAGGCAAGC | *Bam*HI or *Xho*I sites are underlined. |
| | | Nanobody R | ACTGCCACCGCCGCCGTTAACGCTGCTAACGGTAACCTGGGT | |
| | | SNAP F | AACGGCGGCGGTGGCAGTGACAAAGACTGCGAAATGAAGCG | |
| | | SNAP R | TCAGCTTCTGCTCACCGGAACCACCACC*CTCGAG*ACCCAGCCCAGGCTTGCCCA | |

**Table S7.** Protein coding sequences (CDSs) and noncommercial vector used in this work.

| Purpose | Name | CDS or vector sequence | Subcloning note |
|---|---|---|---|
| *E. coli* expression | *Dr*BphP-Avi-His | ATGAGTCGTGACCCTTTGCCATTCTTTCCTCCTCTTTATCTGGGTGGACCCGAGAT<br>TACAACAGAAAACTGCGAACGCGAACCAATTCACATCCCGGGATCTATTCAACCAC<br>ACGGTGCATTGCTGACGGCAGACGGACATTCCGGAGAGGTTTTACAGATGTCGCTT<br>AACGCAGCAACGTTTCTGGGACAAGAGCCTACGGTTTTGCGCGGCCAGACGTTAGC<br>GGCTCTGTTGCCAGAGCAATGGCCGGCCTTACAGGCGGCATTGCCTCCAGGGTGCC<br>CCGATGCATTGCAATACCGCGCGACACTGGATTGGCCGGCGGCAGGACATCTTTCT<br>CTGACAGTCCACCGCGTGGGCGAGCTGTTGATCCTGGAGTTTGAACCTACGGAGGC<br>CTGGGACTCGACTGGCCCGCACGCGTTACGCAATGCGATGTTCGCTCTTGAATCAG<br>CGCCAAACTTGCGCGCGTTAGCTGAAGTGGCCACACAAACCGTACGCGAGCTTACA<br>GGCTTTGACCGCGTGATGTTATACAAATTCGCACCCGATGCGACAGGCGAGGTAAT<br>CGCCGAAGCCCGCCGCGAGGGGTTGCATGCCTTTCTTGGCCATCGTTTTCCGGCCT<br>CAGATATTCCCGCCCAAGCGCGCGCCCTTTACACTCGCCATCTGCTTCGTTTGACT<br>GCGGACACGCGCGCGGCGGCCGTTCCCTTAGACCCAGTACTTAATCCTCAGACTAA<br>CGCTCCTACCCCCTTAGGGGGGGCAGTGCTGCGTGCGACGTCGCCTATGCACATGC<br>AGTACCTTCGCAATATGGGCGTCGGCTCCTCTTTAAGTGTATCAGTGGTAGTTGGG<br>GGGCAGTTATGGGGTCTGATTGCGTGCCATCATCAGACCCCCTATGTTTTGCCACC<br>AGACCTTCGTACTACTCTTGAATACTTGGGGCGTTTATTAAGCCTTCAGGTGCAAG<br>TCAAGGAAGCCGCGGACGTTGCTGCATTCCGTCAGTCACTTCGCGAACACCATGCG<br>CGCGTCGCCTTAGCGGCAGCGCATTCCCTGTCGCCGCACGATACTCTTTCCGACCC<br>TGCACTTGATCTTCTGGGTCTGATGCGTGCTGGGGGCTTAATCCTGCGTTTTGAAG<br>GTCGTTGGCAGACGTTAGGAGAAGTCCCGCCCGCTCCCGCAGTCGATGCACTGCTT<br>GCATGGCTTGAAACCCAACCAGGGGCGCTTGTTCAGACTGATGCATTGGGGCAGTT<br>GTGGCCGGCGGGGGCTGATTTGGCTCCCTCAGCCGCGGGTCTGCTTGCCATTTCAG<br>TAGGGGAGGGATGGAGTGAGTGCTTGGTTTGGTTACGTCCCGAACTGCGCCTTGAG<br>GTTGCGTGGGGTGGAGCAACTCCAGACCAGGCCAAGGACGACCTGGGCCCTCGTCA<br>CAGTTTCGATACTTACTTAGAAGAGAAGCGTGGGTATGCAGAACCCTGGCATCCCG<br>GAGAGATTGAGGAAGCTCAGGATTTGCGCGACACTCTTACTGGCGCATTAAAGCTT<br>GGTGGCGGTAGCGAGAATTTGTATTTTCAGGGTGGCGGTGGCAGTAGCTTATCCAC<br>CCCGCCGACCCCGAGCACTCCTCCTACCGGTCTGAACGACATCTTCGAGGCTCAGA<br>AAATCGAATGGCACGAACATCATCACCACCATCAC | The CDS was inserted into pBAD (Addgene #80341) using *BamHI*/*EcoRI* restriction sites. |
| | *Dr*BphP-His | (*Dr*BphP)-GAATTCCATCATCACCATCACCAT | The sequence of *Dr*BphP is the same as above. The CDS was inserted into pBAD (Addgene #80341) using *BamHI*/*EcoRI* restriction sites. |
| | LDB-3-His | GAAGTTCAGCTGCAGGCAAGCGGTGGTGGTTTTGTTCAGCCTGGTGGTAGCCTGCG<br>TCTGAGCTGTGCAGCCAGCGGTTTTACCTGGGATCATTACATCATGGGCTGGTTTC<br>GCCAGGCACCGGGTAAAGAACGTGAATTTGTTAGCGCAATCAGCGAAAATGGTGAT<br>GCATGGAATTATTATGCCGATAGCGTGAAAGGTCGCTTTACCATTAGCCGTGATAA<br>TAGCAAAAATACCGTTTACCTGCAGATGAATAGTCTGCGTGCAGAAGATACCGCAA<br>CCTATTATTGTGCAATCGGTTTTGATGTTCCATCTGGTCGTTCTTGGCAGGGTTCT<br>CATTTTTGGATGTATTGGGGTCAGGGCACCCAGGTTACCGTTAGCAGCAGCCCGGG<br>AGGCCAACACCATCACCACCATCAT | The CDS was inserted into pADL-23c (Antibody Design Labs) using a *Bgl*I restriction site. |
| | LDB-3-Avi-His | (LDB-3)-<br>AGCCCGGGAGGCCAAAGCTTATCCACCCCGAGTGTAGATCTCGGTGGTCGCCGTAT<br>CATTGGTCTGAACGACATCTTCGAGGCTCAGAAAATCGAATGGCACGAACATCATC<br>ACCACCATCACTCT | The sequence of LDB-3 is the same as above. The CDS was inserted into pADL-23c using a *Bgl*I restriction site. |
| | LDB-14-His | GAAGTTCAGCTGCAGGCAAGCGGTGGTGGTTTTGTTCAGCCTGGTGGTAGCCTGCG<br>TCTGAGCTGTGCAGCCAGCGGTACCACCTCTCGTTGGGAATCTATGGGCTGGTTTC<br>GCCAGGCACCGGGTAAAGAACGTGAATTTGTTAGCGCAATCAGCTGGCAGAATAAT<br>TCTGTTCCATATTATGCCGATAGCGTGAAAGGTCGCTTTACCATTAGCCGTGATAA<br>TAGCAAAAATACCGTTTACCTGCAGATGAATAGTCTGCGTGCAGAAGATACCGCAA<br>CCTATTATTGTGCAGCACAGCATAACTTTCTGGGTCATCGTTATTGGGGTCAGGGC<br>ACCCAGGTTACCGTTAGCAGCAGCCCGGGAGGCCAACACCATCACCACCATCAT | The CDS was inserted into pADL-23c using a *Bgl*I restriction site. |
| | LDB-14-Avi-His | (LDB-14)-<br>AGCCCGGGAGGCCAAAGCTTATCCACCCCGAGTGTAGATCTCGGTGGTCGCCGTAT<br>CATTGGTCTGAACGACATCTTCGAGGCTCAGAAAATCGAATGGCACGAACATCATC<br>ACCACCATCACTCT | The sequence of LDB-14 is the same as above. The CDS was inserted into pADL-23c using a *Bgl*I restriction site. |

| | | | |
|---|---|---|---|
| Yeast two-hybrid | GAL4-*Dr*BphP | ATGAAGCTACTGTCTTCTATCGAACAAGCATGCGATATTTGCCGACTTAAAAAGCT<br>CAAGTGCTCCAAAGAAAAACCGAAGTGCGCCAAGTGTCTGAAGAACAACTGGGAGT<br>GTCGCTACTCTCCCAAAACCAAAAGGTCTCCGCTGACTAGGGCACATCTGACAGAA<br>GTGGAATCAAGGCTAGAAAGACTGGAACAGCTATTTCTACTGATTTTTCCTCGAGA<br>AGACCTTGACATGATTTTGAAAATGGATTCTTTACAGGATATAAAAGCATTGTTAA<br>CAGGATTATTTGTACAAGATAATGTGAATAAAGATGCCGTCACAGATAGATTGGCT<br>TCAGTGGAGACTGATATGCCTCTAACATTGAGACAGCATAGAATAAGTGCGACATC<br>ATCATCGGAAGAGAGTAGTAACAAAGGTCAAAGACAGTTGACTGTATCGCCGGAAT<br>TTGTAATACGACTCACTATAGGGCGAGCCGCCATCATGGAGGAGCAGAAGCTGATC<br>TCAGAGGAGGACCTGCAT-(*Dr*BphP) | The sequence of *Dr*BphP is the same as above. The CDS was inserted into pGBKT7 vector (Clontech) using *Nde*I/*BamH*I restriction sites. |
| | AD-LDB-3 | ATGGATAAAGCGGAATTAATTCCCGAGCCTCCAAAAAAGAAGAGAAAGGTCGAATT<br>GGGTACCGCCGCCAATTTTAATCAAAGTGGGAATATTGCTGATAGCTCATTGTCCT<br>TCACTTTCACTAACAGTAGCAACGGTCCGAACCTCATAACAACTCAAACAAATTCT<br>CAAGCGCTTTCACAACCAATTGCCTCCTCTAACGTTCATGATAACTTCATGAATAA<br>TGAAATCACGGCTAGTAAAATTGATGATGGTAATAATTCAAAACCACTGTCACCTG<br>GTTGGACGGACCAAACTGCGTATAACGCGTTTGGAATCACTACAGGGATGTTTAAT<br>ACCACTACAATGGATGATGTATATAACTATCTATTCGATGATGAAGATACCCCACC<br>AAACCCAAAAAAAGAGATCTTTAATACGACTCACTATAGGGCGAGCGCCGCCATGG<br>AGTACCCATACGACGTACCAGATTACGCTCATATGGGAGGCGGTTCCGGTGGCGGT<br>TCT-(LDB-3) | The sequence of LDB-3 is the same as above. The CDS was inserted into pGADT7 (Clontech) using *Nde*I/*BamH*I restriction sites. |
| | AD-LDB-4 | (AD)-<br>ATCTTTAATACGACTCACTATAGGGCGAGCGCCGCCATGGAGTACCCATACGACGT<br>ACCAGATTACGCTCATATGGGAGGCGGTTCCGGTGGCGGTTCTGAAGTTCAGCTGC<br>AGGCAAGCGGTGGTGGTTTTGTTCAGCCTGGTGGTAGCCTGCGTCTGAGCTGTGCA<br>GCCAGCGGTGATACCTCTTACCTGTACTCTATGGGCTGGTTTCGCCAGGCACCGGG<br>TAAAGAACGTGAATTTGTTAGCGCAATCAGCTGGTGGTGGAATCTGACTCAGTATT<br>ATGCCGATAGCGTGAAAGGTCGCTTTACCATTAGCCGTGATAATAGCAAAAATACC<br>GTTTACCTGCAGATGAATAGTCTGCGTGCAGAAGATACCGCAACCTATTATTGTGC<br>ATGGTCTATCTACTTTCCACCAGGTAACGATTACAACGGTTACCATTATTGGGGTC<br>AGGGCACCCAGGTTACCGTTAGCAGC | The CDS was inserted into pGADT7 using *Nde*I/*BamH*I restriction sites. The sequences of AD and LDB-14 are the same as above. |
| | AD-LDB-6 | (AD)-<br>ATCTTTAATACGACTCACTATAGGGCGAGCGCCGCCATGGAGTACCCATACGACGT<br>ACCAGATTACGCTCATATGGGAGGCGGTTCCGGTGGCGGTTCTGAAGTTCAGCTGC<br>AGGCAAGCGGTGGTGGTTTTGTTCAGCCTGGTGGTAGCCTGCGTCTGAGCTGTGCA<br>GCCAGCGGTTTTTTTTCTAACTGGTCTGATATGGGCTGGTTTCGCCAGGCACCGGG<br>TAAAGAACGTGAATTTGTTAGCGCAATCAGCTTTTGGGCAGATGGTACTGAATATT<br>ATGCCGATAGCGTGAAAGGTCGCTTTACCATTAGCCGTGATAATAGCAAAAATACC<br>GTTTACCTGCAGATGAATAGTCTGCGTGCAGAAGATACCGCAACCTATTATTGTGC<br>ATGGTACGGTCCAGTTAACGGTTTTTACATGTTTGATTATTGGGGTCAGGGCACCC<br>AGGTTACCGTTAGCAGC | |
| | AD-LDB-7 | (AD)-<br>ATCTTTAATACGACTCACTATAGGGCGAGCGCCGCCATGGAGTACCCATACGACGT<br>ACCAGATTACGCTCATATGGGAGGCGGTTCCGGTGGCGGTTCTGAAGTTCAGCTGC<br>AGGCAAGCGGTGGTGGTTTTGTTCAGCCTGGTGGTAGCCTGCGTCTGAGCTGTGCA<br>GCCAGCGGTTCTACCTCTGATTTTGAATCTATGGGCTGGTTTCGCCAGGCACCGGG<br>TAAAGAACGTGAATTTGTTAGCGCAATCAGCTCTTGGTTTACTAATCCACCATATT<br>ATGCCGATAGCGTGAAAGGTCGCTTTACCATTAGCCGTGATAATAGCAAAAATACC<br>GTTTACCTGCAGATGAATAGTCTGCGTGCAGAAGATACCGCAACCTATTATTGTGC<br>ACATCGTTCTATCTGGTACCATCCAACCTATTGGGGTCAGGGCACCCAGGTTACCG<br>TTAGCAGC | |
| | AD-LDB-14 | (AD)-<br>ATCTTTAATACGACTCACTATAGGGCGAGCGCCGCCATGGAGTACCCATACGACGT<br>ACCAGATTACGCTCATATGGGAGGCGGTTCCGGTGGCGGTTCT-(LDB-14) | |
| Mammalian two-hybrid | GAL4-*Dr*BphP | GAL4-*Dr*BphP | The sequence of GAL4-*Dr*BphP is the same as above. The CDS was inserted into pBobi (see below for the sequence) using *BamH*I/*Xho*I restriction sites. |

| | | | |
|---|---|---|---|
| | NLS-<br>LDB-3-<br>p65 | ATGGGATCCCCCAAGAAGAAGCGCAAGGTGGAAGCTAGCGCTTCCCCGAAGAAAAA<br>GCGGAAAGTCGAGGCCTCCGCATCTCCAAAAAAAAAAAGCAAGGTTGAAGCATCTG<br>GATCCGGTACCGGAGGAAGTGGCAGCTCTGGCGGCAGTGGAGGGTCTGGTGGCAGC<br>GGA-(LDB-3)-<br>AGCGGCGGCGGTGGCAGTCAGTACCTGCCAGATACAGACGATCGTCACCGGATTGA<br>GGAGAAACGTAAAAGGACATATGAGACCTTCAAGAGCATCATGAAGAAGAGTCCTT<br>TCAGCGGACCCACCGACCCCCGGCCTCCACCTCGACGCATTGCTGTGCCTTCCCGC<br>AGCTCAGCTTCTGTCCCCAAGCCAGCACCCCAGCCCTATCCCTTTACGTCATCCCT<br>GAGCACCATCAACTATGATGAGTTTCCCACCATGGTGTTTCCTTCTGGGCAGATCA<br>GCCAGGCCTCGGCCTTGGCCCCGGCCCCTCCCCAAGTCCTGCCCCAGGCTCCAGCC<br>CCTGCCCCTGCTCCAGCCATGGTATCAGCTCTGGCCCAGGCCCCAGCCCCTGTCCC<br>AGTCCTAGCCCCAGGCCCTCCTCAGGCTGTGGCCCCACCTGCCCCCAAGCCCACCC<br>AGGCTGGGGAAGGAACGCTGTCAGAGGCCCTGCTGCAGCTGCAGTTTGATGATGAA<br>GACCTGGGGGCCTTGCTTGGCAACAGCACAGACCCAGCTGTGTTCACAGACCTGGC<br>ATCCGTCGACAACTCCGAGTTTCAGCAGCTGCTGAACCAGGGCATACCTGTGGCCC<br>CCCACACAACTGAGCCCATGCTGATGGAGTACCCTGAGGCTATAACTCGCCTAGTG<br>ACAGGGGCCCAGAGGCCCCCCGACCCAGCTCCTGCTCCACTGGGGGCCCCGGGGCT<br>CCCCAATGGCCTCCTTTCAGGAGATGAAGACTTCTCCTCCATTGCGGACATGGACT<br>TCTCAGCCCTGCTGAGTCAGATCAGCTCC | The CDS was inserted into pBobi vector using *BamH*I/*Xho*I restriction sites. The sequence of LDB-3 is the same as above. |
| | NLS-<br>LDB-4-<br>p65 | (NLS)-<br>GGATCCGGTACCGGAGGAAGTGGCAGCTCTGGCGGCAGTGGAGGGTCTGGTGGCAG<br>CGGA-(LDB-4)-AGCGGCGGCGGTGGCAGT-(p65) | The sequences of NLS, nanobody and p65 are the same as above. The CDS was inserted into pBobi using *BamH*I/*Xho*I restriction sites. |
| | NLS-<br>LDB-6-<br>p65 | (NLS)-<br>GGATCCGGTACCGGAGGAAGTGGCAGCTCTGGCGGCAGTGGAGGGTCTGGTGGCAG<br>CGGA-(LDB-6)-AGCGGCGGCGGTGGCAGT-(p65) | |
| | NLS-<br>LDB-7-<br>p65 | (NLS)-<br>GGATCCGGTACCGGAGGAAGTGGCAGCTCTGGCGGCAGTGGAGGGTCTGGTGGCAG<br>CGGA-(LDB-7)-AGCGGCGGCGGTGGCAGT-(p65) | |
| | NLS-<br>LDB-<br>14-p65 | ATGCCCAAGAAGAAGCGCAAGGTGGAAGCTAGCGCTTCCCCGAAGAAAAAGCGGAA<br>AGTCGAGGCCTCCGCATCTCCAAAAAAAAAAAGCAAGGTTGAAGCATCTGGATCCG<br>GAGGCGGTTCCGGTGGCGGTTCT-(LDB-14)-<br>GGTACCGGAGGAAGTGGCAGCTCTGGCGGCAGTGGAGGGTCTGGTGGCAGCGGA-<br>(p65)-GGAGGAAGTGGCAGCTCTGGCGGCAGTGGA | The CDS was inserted into pcDNA3 (Invitrogen) using *Hind*III/*Xho*I restriction sites. The sequences of LDB-14 and p65 are the same as above. |
| | GAL4-<br>*Rp*Bph<br>P1 | (GAL4)-<br>CCGGAATTTGTAATACGACTCACTATAGGGCGAGCCGCCATCATGGAGGAGCAGAA<br>GCTGATCTCAGAGGAGGACCTGCATGTGGCAGGTCATGCCTCTGGCAGCCCCGCAT<br>TCGGGACCGCCGATCTTTCGAATTGCGAACGTGAAGAGATCCACCTCGCCGGCTCG<br>ATCCAGCCGCATGGCGCGCTTCTGGTCGTCAGCGAGCCGGATCATCGCATCATCCA<br>GGCCAGCGCCAACGCCGCGGAATTTCTGAATCTCGGAAGCGTGCTCGGCGTTCCGC<br>TCGCCGAGATCGACGGCGATCTGTTGATCAAGATCCTGCCGCATCTCGATCCCACC<br>GCCGAAGGCATGCCGGTCGCGGTGCGCTGCCGGATCGGCAATCCCTCCACGGAGTA<br>CGACGGTCTGATGCATCGGCCTCCGGAAGGCGGGCTGATCATCGAGCTCGAACGTG<br>CCGGCCCGCCGATCGATCTGTCCGGCACGCTGGCGCCGGCGCTGGAGCGGATCCGC<br>ACGGCGGGCTCGCTGCGCGCGCTGTGCGATGACACCGCGCTGCTGTTTCAGCAGTG<br>CACCGGCTACGACCGGGTGATGGTGTATCGCTTCGACGAGCAGGGCCACGGCGAAG<br>TGTTCTCCGAGCGCCACGTGCCCGGGCTCGAATCCTATTTCGGCAACCGCTATCCG<br>TCGTCGGACATTCCGCAGATGGCGCGGCGGCTGTACGAGCGGCAGCGCGTCCGCGT<br>GCTGGTCGACGTCAGCTATCAGCCGGTGCCGCTGGAGCCGCGGCTGTCGCCGCTGA<br>CCGGGCGCGATCTCGACATGTCGGGCTGCTTCCTGCGCTCGATGTCGCCGATCCAT<br>CTGCAGTACCTGAAGAACATGGGCGTGCGCGCCACCCTGGTGGTGTCGCTGGTGGT<br>CGGCGGCAAGCTGTGGGGCCTGGTTGCCTGTCACCATTATCTGCCGCGCTTCATCC<br>ATTTCGAGCTGCGGGCGATCTGCGAACTGCTCGCCGAAGCGATCGCGACGCGGATC<br>ACCGCGCTTGAGAGCTTCGCGCAGAGCCAGTCGGAGCTGTTCGTGCAGCGGCTCGA<br>ACAGCGCATGATCGAAGCGATCACCCGTGAAGGCGATTGGCGCGCAGCGATTTTCG<br>ACACCAGCCAATCGATCCTGCAGCCGCTGCACGCCGACGGTTGCGCGCTGGTGTAC<br>GAAGACCAGATCAGGACCATCGGTGACGTACCTTCCACGCAGGATGTTCGCGAGAT<br>CGCCGGGTGGCTCGATCGCCAGCCACGTGCGGCGGTGACCTCGACCGCGTCGCTCG<br>GTCTCGACGTGCCGGAGCTCGCGCATCTGACGCGGATGGCGAGCGGCGTGGTCGCG<br>GCGCCGATTTCGGATCATCGCGGCGAGTTTCTGATGTGGTTCCGCCCCGAGCGCGT<br>CCACACCGTTACCTGGGGCGGCGATCCGAAGAAGCCGTTCACGATGGGCGATACAC<br>CGGCGGATCTGTCGCCGCGGCGCTCCTTCGCCAAATGGCATCAGGTTGTCGAAGGC<br>ACGTCCGATCCGTGGACGGCCGCCGATCTCGCCGCGGCTCGCACCATCGGTCAGAC<br>CGTCGCCGACATCGTGCTGCAATTCCGCGCGGTGCGGACACTGATCGCCCGCGAAC<br>AGTACGAACAGTTTTCGTCCCAGGTGCACGCTTCGATGCAGCCGGTGCTGATCACC<br>GACGCCGAAGGCCGCATCCTGCTGATGAACGACTCGTTCCGCGACATGTTGCCGGC<br>GGGGTCGCCATCCGCCGTCCATCTCGACGATCTCGCCGGGTTCTTCGTCGAATCGA<br>ACGATTTCCTGCGCAACGTCGCCGAACTGATCGATCACGGCCGCGGGTGGCGCGGC | The CDS was subcloned into pcDNA3 using *Hind*III/*Xho*I restriction sites. The sequence of GAL4 is the same as above. |

| | | | |
|---|---|---|---|
| | | GAAGTTCTGCTGCGCGGCGCAGGTAATCGCCCGTTGCCGCTGGCAGTGCGCGCCGA<br>TCCGGTGACGCGCACGGAGGACCAGTCGCTCGGCTTCGTGCTGATCTTCAGCGACG<br>CTACCGATCGTCGCACCGCAGATGCCGCACGCACGCGTTTCCAGGAAGGCATTCTT<br>GCCAGCGCACGTCCCGGCGTGCGGCTCGACTCCAAGTCCGACCTCTTGCACGAGAA<br>GCTGCTGTCCGCGCTGGTCGAGAACGCGCAGCTTGCCGCATTGGAAATTACTTACG<br>GCGTCGAGACCGGACGCATCGCCGAGCTGCTCGAAGGCGTTCGCCAGTCGATGCTG<br>CGCACCGCCGAAGTGCTCGGCCATCTGGTGCAGCACGCGGCGCGCACGGCCGGCAG<br>CGACAGCTCGAGCAATGGCTCGCAGAACAAGAAG | |
| | NLS-PpsR2-p65 | ATGCCCAAGAAGAAGCGCAAGGTGGAAGCTAGCGCTTCCCCGAAGAAAAAGCGGAA<br>AGTCGAGGCCTCCGCATCTCCAAAAAAAAAAAGCAAGGTTGAAGCATCTGGATCCG<br>GAGGCGGTTCCGGTGGCGGTTCTGTGGCGTCAAAGTCCGTTCATGCCGACATCACC<br>CTTCTGCTCGATATGGAGGGTGTGATTCGCGAAGCCACCCTGTCTCCGACGATGGC<br>GGCCGAGAGCGTGGACGGTTGGCTGGGGCGTCGCTGGAGCGACATCGCCGGCGCCG<br>AAGGCGGCGACAAGGTTCGCCGCATGGTCGAAGACGCCCGCCGCAGCGGCATCTCG<br>GCTTTCCGCCAGATCAATCAGCCTTTCCCGAGCGGCGTCGAAATCCCGATCGAATT<br>CACCACGATGCTGCTGGGCGACCGCACCGGCATGATCGCGGTCGGCAAGAACATGC<br>AGGCGGTCACCGAGCTGCATTCCCGGCTGATCGCTGCGCAGCAGGCGATGGAGCGC<br>GACTATTGGCGGTTGCGTGAATTGGAGACTCGCTACCGCCTGGTGTTCGACGCTGC<br>CGCCGATGCGGTGATGATCGTCTCCGCCGGCGACATGCGCATCGTCGAAGCCAACC<br>GGGCGGCGGTGAATGCGATCAGCCGCGTCGAGCGCGGCAATGACGACCTTGCGGGG<br>CGTGATTTCCTCGCCGAAGTGGCGGCTGCCGATCGCGATGCGGTGCGCGACATGCT<br>GGCCCAGGTGCGTCAGCGCGGCACCGCACTCAGCGTCCTCGTTCATCTCGGCCGTT<br>ACGACCGCGCCTGGATGCTGCGCGGTTCGCTGATGTCGTCCGAGCGTCGTCAGGTT<br>TTCCTGCTGCACTTCACCCCGGTGACCACGACTCCCGCGATCGACGACGTCGACGA<br>TGATGCCGTGCTGCGCGGGCTGATCGATCGCATTCCCGACGGGTTCGTCGCACTGG<br>ATTCGGAAGGCGTCGTTCGTCACGCCAACCAGGCGTTTCTCGATCTGGTCCAGATC<br>GGCTCCAAGCCTGCGGCGGTCGGACGATCGCTGGGCGTCTGGATGGGTCGTCCGGG<br>CGCCGATCTGTCCAGCTTGCTGACGCTGCTGCGGCGCTACAAGACGGTGCGGCTGT<br>TCCAAACGACGATCCGCGGCGAGCTCGGCACCGAGACTGAAGTCGAGGTCTCGGCC<br>GTCGACGGCGAGGACGACCAATACATCGGCGTTCTGATGCGCAATGTCGCGCGACG<br>CCTCGACGCTGCGGACGACCACGATGCCTTGCGTCAGGCGCTCGGCCCGATCAGCA<br>AGCAGCTCGGGCGATCCTCGCTGCGCAAGCTGGTGAAGAACGCCGTGAGCATTGTC<br>GAGCAGCACTACGTGAAGGAAGCGCTGTTGCGATCCAAGGGCAATCGCACGGCAAC<br>TGCCGAACTGCTCGGATTGAGCCGGCAGAGCCTTTATGCAAAACTCAACTCCTACG<br>GCTTCGACGACAAAGGTGTCGTTGCTTCTGCTGCCGACGGTGCAGAGGGCGCCTCA<br>GACGACGCAGAGGATGGTACCGGAGGAAGTGGCAGCTCTGGCGGCAGTGGAGGGTC<br>TGGTGGCAGCGGA-(p65)-GGAGGAAGTGGCAGCTCTGGCGGCAGTGGA | The CDSs were inserted into pcDNA3 using *Hind*III/*Xho*I restriction sites. The sequence of p65 is the same as above. |
| | NLS-Q-PAS1-p65 | ATGCCCAAGAAGAAGCGCAAGGTGGAAGCTAGCGCTTCCCCGAAGAAAAAGCGGAA<br>AGTCGAGGCCTCCGCATCTCCAAAAAAAAAAAGCAAGGTTGAAGCATCTGGATCCG<br>GAGGCGGTTCCGGTGGCGGTTCTGGCAAGAACATGCAGGCGGTCACCGAGCTGCAT<br>TCCCGGCTGATCGCTGCGCAGCAGGCGATGGAGCGCGACTATTGGCGGTTGCGTGA<br>ATTGGAGACTCGCTACCGCCTGGTGTTCGACGCTGCCGCCGATGCGGTGATGATCG<br>TCTCCGCCGGCGACATGCGCATCGTCGAAGCCAACCGGGCGGCGGTGAATGCGATC<br>AGCCGCGTCGAGCGCGGCAATGACGACCTTGCGGGGCGTGATTTCCTCGCCGAAGT<br>GGCGGCTGCCGATCGCGATGCGGTGCGCGACATGCTGGCCCAGGTGCGTCAGCGCG<br>GCACCGCACTCAGCGTCCTCGTTCATCTCGGCCGTTACGACCGCGCCTGGATGCTG<br>CGCGGTTCGCTGATGTCGTCCGAGCGTCGTCAGGTTTTCCTGCTGCACTTCACCCC<br>GGTGACCACGACTCCCGCGATCGACGACGGTACCGGAGGAAGTGGCAGCTCTGGCG<br>GCAGTGGAGGGTCTGGTGGCAGCGGA-(p65)-<br>GGAGGAAGTGGCAGCTCTGGCGGCAGTGGA | |
| Detection of nanobody expression in mammalian cells | LDB-3-SNAP | ATGGGATCC-(LDB-3)-<br>GTTAACGGCGGCGGTGGCAGTGACAAAGACTGCGAAATGAAGCGCACCACCCTGGA<br>TAGCCCTCTGGGCAAGCTGGAACTGTCTGGGTGCGAACAGGGCCTGCACCGTATCA<br>TCTTCCTGGGCAAAGGAACATCTGCCGCCGACGCCGTGGAAGTGCCTGCCCCAGCC<br>GCCGTGCTGGGCGGACCAGAGCCACTGATGCAGGCCACCGCCTGGCTCAACGCCTA<br>CTTTCACCAGCCTGAGGCCATCGAGGAGTTCCCTGTGCCAGCCCTGCACCACCCAG<br>TGTTCCAGCAGGAGAGCTTTACCCGCCAGGTGCTGTGGAAACTGCTGAAAGTGGTG<br>AAGTTCGGAGAGGTCATCAGCTACAGCCACCTGGCCGCCCTGGCCGGCAATCCCGC<br>CGCCACCGCCGCCGTGAAAACCGCCCTGAGCGGAAATCCCGTGCCCATTCTGATCC<br>CCTGCCACCGGGTGGTGCAGGGCGACCTGGACGTGGGGGGCTACGAGGGCGGGCTC<br>GCCGTGAAAGAGTGGCTGCTGGCCCACGAGGGCCACAGACTGGGCAAGCCTGGGCT<br>GGGT | The CDS was inserted into pBobi vector using *BamH*I/*Xho*I restriction sites. The sequence of LDB-3 is the same as above. |
| | LDB-4-SNAP | ATGGGATCC-(LDB-4)-GTTAACGGCGGCGGTGGCAGT-(SNAP) | The CDSs were inserted into pBobi using *BamH*I/*Xho*I restriction sites. The sequences of nanobodies and |
| | LDB-6-SNAP | ATGGGATCC-(LDB-6)-GTTAACGGCGGCGGTGGCAGT-(SNAP) | |
| | LDB-7-SNAP | ATGGGATCC-(LDB-7)-GTTAACGGCGGCGGTGGCAGT-(SNAP) | |

| | | | |
|---|---|---|---|
| | LDB-14-SNAP | ATGGGATCC-(LDB-14)-GTTAACGGCGGCGGTGGCAGT-(SNAP) | SNAP are the same as above. |
| | pBobi vector | TGACGGATCGGGAGATCTCCCGATCCCCTATGGTCGACTCTCAGTACAATCTGCTC<br>TGATGCCGCATAGTTAAGCCAGTATCTGCTCCCTGCTTGTGTGTTGGAGGTCGCTG<br>AGTAGTGCGCGAGCAAAATTTAAGCTACAACAAGGCAAGGCTTGACCGACAATTGC<br>ATGAAGAATCTGCTTAGGGTTAGGCGTTTTGCGCTGCTTCGCGATGTACGGGCCAG<br>ATATACGCGTTGACATTGATTATTGACTAGTTATTAATAGTAATCAATTACGGGGT<br>CATTAGTTCATAGCCCATATATGGAGTTCCGCGTTACATAACTTACGGTAAATGGC<br>CCGCCTGGCTGACCGCCCAACGACCCCCGCCCATTGACGTCAATAATGACGTATGT<br>TCCCATAGTAACGCCAATAGGGACTTTCCATTGACGTCAATGGGTGGACTATTTAC<br>GGTAAACTGCCCACTTGGCAGTACATCAAGTGTATCATATGCCAAGTACGCCCCCT<br>ATTGACGTCAATGACGGTAAATGGCCCGCCTGGCATTATGCCCAGTACATGACCTT<br>ATGGGACTTTCCTACTTGGCAGTACATCTACGTATTAGTCATCGCTATTACCATGG<br>TGATGCGGTTTTGGCAGTACATCAATGGGCGTGGATAGCGGTTTGACTCACGGGGA<br>TTTCCAAGTCTCCACCCCATTGACGTCAATGGGAGTTTGTTTTGGCACCAAAATCA<br>ACGGGACTTTCCAAAATGTCGTAACAACTCCGCCCCATTGACGCAAATGGGCGGTA<br>GGCGTGTACGGTGGGAGGTCTATATAAGCAGCGCGTTTTGCCTGTACTGGGTCTCT<br>CTGGTTAGACCAGATCTGAGCCTGGGAGCTCTCTGGCTAACTAGGGAACCCACTGC<br>TTAAGCCTCAATAAAGCTTGCCTTGAGTGCTTCAAGTAGTGTGTGCCCGTCTGTTG<br>TGTGACTCTGGTAACTAGAGATCCCTCAGACCCTTTTAGTCAGTGTGGAAAATCTC<br>TAGCAGTGGCGCCCGAACAGGGACCTGAAAGCGAAAGGGAAACCAGAGCTCTCTCG<br>ACGCAGGACTCGGCTTGCTGAAGCGCGCACGGCAAGAGGCGAGGGGCGGCGACTGG<br>TGAGTACGCCAAAAATTTTGACTAGCGGAGGCTAGAAGGAGAGAGATGGGTGCGAG<br>AGCGTCAGTATTAAGCGGGGGAGAATTAGATCGCGATGGGAAAAAATTCGGTTAAG<br>GCCAGGGGGAAAGAAAAAATATAAATTAAAACATATAGTATGGGCAAGCAGGGAGC<br>TAGAACGATTCGCAGTTAATCCTGGCCTGTTAGAAACATCAGAAGGCTGTAGACAA<br>ATACTGGGACAGCTACAACCATCCCTTCAGACAGGATCAGAAGAACTTAGATCATT<br>ATATAATACAGTAGCAACCCTCTATTGTGTGCATCAAAGGATAGAGATAAAAGACA<br>CCAAGGAAGCTTTAGACAAGATAGAGGAAGAGCAAAACAAAAGTAAGACCACCGCA<br>CAGCAAGCGGCCGCTGATCTTCAGACTTGGAGGAGGAGATATGAGGGACAATTGGA<br>GAAGTGAATTATATAAATATAAAGTAGTAAAAATTGAACCATTAGGAGTAGCACCC<br>ACCAAGGCAAAGAGAAGAGTGGTGCAGAGAGAAAAAAGAGCAGTGGGAATAGGAGC<br>TTTGTTCCTTGGGTTCTTGGGAGCAGCAGGAAGCACTATGGGCGCAGCCTCAATGA<br>CGCTGACGGTACAGGCCAGACAATTATTGTCTGGTATAGTGCAGCAGCAGAACAAT<br>TTGCTGAGGGCTATTGAGGCGCAACAGCATCTGTTGCAACTCACAGTCTGGGGCAT<br>CAAGCAGCTCCAAGCAAGAATCCTAGCTGTGGAAAGATACCTAAAGGATCAACAGC<br>TCCTAGGGATTTGGGGTTGCTCTGGAAAACTCATTTGCACCACTGCTGTGCCTTGG<br>AATGCTAGTTGGAGTAATAAATCTCTGGAACAGATCTGGAATCACACGACCTGGAT<br>GGAGTGGGACAGAGAAATTAACAATTACACAAGCTTAATACACTCCTTAATTGAAG<br>AATCGCAAAACCAGCAAGAAAAGAATGAACAAGAATTATTGGAATTAGATAAATGG<br>GCAAGTTTGTGGAATTGGTTTAACATAACAAATTGGCTGTGGTATATAAAATTATT<br>CATAATGATAGTAGGAGGCTTGGTAGGTTTAAGAATAGTTTTTGCTGTACTTTCTA<br>TAGTGAATAGAGTTAGGCAGGGATATTCACCATTATCGTTTCAGACCCACCTCCCA<br>ATCCCGAGGGGACCCGACAGGCCCGAAGGAATAGAAGAAGAAGGTGGAGAGAGAGA<br>CAGAGACAGATCCATTCGATTAGTGAACGGATCAACTTTTAAAAGAAAAGGGGGGA<br>TTGGGGGGTACAGTGCAGGGGAAAGAATAGTAGACATAATAGCAACAGACATACAA<br>ACTAAAGAATTACAAAAACAAATTACAAAAATTCAAAATTTTATCGATAAGCTTGG<br>GAGTTCCGCGTTACATAACTTACGGTAAATGGCCCGCCTGGCTGACCGCCCAACGA<br>CCCCCGCCCATTGACGTCAATAATGACGTATGTTCCCATAGTAACGCCAATAGGGA<br>CTTTCCATTGACGTCAATGGGTGGAGTATTTACGGTAAACTGCCCACTTGGCAGTA<br>CATCAAGTGTATCATATGCCAAGTACGCCCCCTATTGACGTCAATGACGGTAAATG<br>GCCCGCCTGGCATTATGCCCAGTACATGACCTTATGGGACTTTCCTACTTGGCAGT<br>ACATCTACGTATTAGTCATCGCTATTACCATGGTGATGCGGTTTTGGCAGTACATC<br>AATGGGCGTGGATAGCGGTTTGACTCACGGGGATTTCCAAGTCTCCACCCCATTGA<br>CGTCAATGGGAGTTTGTTTTGGCACCAAAATCAACGGGACTTTCCAAAATGTCGTA<br>ACAACTCCGCCCCATTGACGCAAATGGGCGGTAGGCGTGTACGGTGGGAGGTCTAT<br>ATAAGCAGAGCTCGTTTAGTGAACCGTCAGATCGCCTGGAGACGCCATCCACGCTG<br>TTTTGACCTCCATAGAAGACACCGACTGAGCTCCTTAAGGTTGCCACCATGGGATC<br>CCTCGAGGGTGGTGGTTCCGGTGAGCAGAAGCTGATCTCAGAGGAGGACCTGTGAT<br>CGAGCCATGGAAGCTTGATATCTAACTGACTGAACCGGTGGTACCGAGGAATTAAT<br>TCGCTGTCTGCGAGGGCCAGCTGTTGGGGTGAGTACTCCCTCTCAAAAGCGGGCAT<br>GACTTCTGCGCTAAGATTGTCAGTTTCCAAAAACGAGGAGGATTTGATATTCACCT<br>GGCCCGCGGTGATGCCTTTGAGGGTGGCCGCGTCCATCTGGTCAGAAAAGACAATC<br>TTTTTGTTGTCAAGCTTGAGGTGTGGCAGGCTTGAGATCTGGCCATACACTTGAGT<br>GACAATGACATCCACTTTGCCTTTCTCTCCACAGGTGTCCACTCCCAGGTCCAACT<br>GCAGGTCGAGCATGCATCTAGGGCGGCCAATTCCGCCGATCCTGGAAAAACATGGA<br>GCAATCACAAGTAGCAATACAGCAGCTACCAATGCTGATTGTGCCTGGCTAGAAGC<br>ACAAGAGGAGGAGGAGGTGGGTTTTCCAGTCACACCTCAGACAATCAACCTCTGGA<br>TTACAAAATTTGTGAAAGATTGACTGGTATTCTTAACTATGTTGCTCCTTTTACGC<br>TATGTGGATACGCTGCTTTAATGCCTTTGTATCATGCTATTGCTTCCCGTATGGCT | |

| | | |
|---|---|---|
| | | TTCATTTTCTCCTCCTTGTATAAATCCTGGTTGCTGTCTCTTTATGAGGAGTTGTG<br>GCCCGTTGTCAGGCAACGTGGCGTGGTGTGCACTGTGTTTGCTGACGCAACCCCCA<br>CTGGTTGGGGCATTGCCACCACCTGTCAGCTCCTTTCCGGGACTTTCGCTTTCCCC<br>CTCCCTATTGCCACGGCGGAACTCATCGCCGCCTGCCTTGCCCGCTGCTGGACAGG<br>GGCTCGGCTGTTGGGCACTGACAATTCCGTGGTGTTGTCGGGGAAGCTGACGTCCT<br>TTCCATGGCTGCTCGCCTGTGTTGCCACCTGGATTCTGCGCGGGACGTCCTTCTGC<br>TACGTCCCTTCGGCCCTCAATCCAGCGGACCTTCCTTCCCGCGGCCTGCTGCCGGC<br>TCTGCGGCCTCTTCCGCGTCTTCGCCTTCGCCCTCAGACGAGTCGGATCTCCCTTT<br>GGGCCGCCTCCCCGCCTGGAATTCGAGCTCGGTACCTTTAAGACCAATGACTTACA<br>AGGCAGCTGTAGATCTTAGCCACTTTTTAAAAGAAAAGGGGGGACTGGAAGGGCTA<br>ATTCACTCCCAAAGAAGACAAGATATCCTTGATCTGTGGATCTACCACACACAAGG<br>CTACTTCCCTGATTGACAGAACTACACACCAGGGCCAGGGGTCAGATATCCACTGA<br>CCTTTGGATGGTGCTACAAGCTAGTACCAGTTGAGCCAGATAAGATAGAAGAGGCC<br>AATAAAGGAGAGAACACCAGCTTGTTACACCCTGTGAGCCTGCATGGGATGGATGA<br>CCCGGAGAGAGAAGTGTTAGAGTGGAGGTTTGACAGCCGCCTAGCATTTCATCACG<br>TGGCCCGAGAGCTGCATCCGGACTGTACTGGGTCTCTCTGGTTAGACCAGATCTGA<br>GCCTGGGAGCTCTCTGGCTAACTAGGGAACCCACTGCTTAAGCCTCAATAAAGCTT<br>GCCTTGAGTGCTTCAAGTAGTGTGTGCCCGTCTGTTGTGTGACTCTGGTAACTAGA<br>GATCCCTCAGACCCTTTTAGTCAGTGTGGAAAATCTCTAGCAGGGCCGCTTTAAAC<br>CCGCTGATCAGCCTCGACTGTGCCTTCTAGTTGCCAGCCATCTGTTGTTTGCCCCT<br>CCCCCGTGCCTTCCTTGACCCTGGAAGGTGCCACTCCCACTGTCCTTTCCTAATAA<br>AATGAGGAAATTGCATCGCATTGTCTGAGTAGGTGTCATTCTATTCTGGGGGGTGG<br>GGTGGGGCAGGACAGCAAGGGGGAGGATTGGGAAGACAATAGCAGGCATGCTGGGG<br>ATGCGGTGGGCTCTATGGCTTCTGAGGCGGAAAGAACCAGCTGGGGCTCTAGGGGG<br>TATCCCCACGCGCCCTGTAGCGGCGCATTAAGCGCGGCGGGTGTGGTGGTTACGCG<br>CAGCGTGACCGCTACACTTGCCAGCGCCCTAGCGCCCGCTCCTTTCGCTTTCTTCC<br>CTTCCTTTCTCGCCACGTTCGCCGGCTTTCCCCGTCAAGCTCTAAATCGGGGCATC<br>CCTTTAGGGTTCCGATTTAGTGCTTTACGGCACCTCGACCCCAAAAAACTTGATTA<br>GGGTGATGGTTCACGTAGTGGGCCATCGCCCTGATAGACGGTTTTTCGCCCTTTGA<br>CGTTGGAGTCCACGTTCTTTAATAGTGGACTCTTGTTCCAAACTGGAACAACACTC<br>AACCCTATCTCGGTCTATTCTTTTGATTTATAAGGGATTTTGGGGATTTCGGCCTA<br>TTGGTTAAAAAATGAGCTGATTTAACAAAAATTTAACGCGAATTAATTCTGTGGAA<br>TGTGTGTCAGTTAGGGTGTGGAAAGTCCCCAGGCTCCCCAGGCAGGCAGAAGTATG<br>CAAAGCATGCATCTCAATTAGTCAGCAACCAGGTGTGGAAAGTCCCCAGGCTCCCC<br>AGCAGGCAGAAGTATGCAAAGCATGCATCTCAATTAGTCAGCAACCATAGTCCCGC<br>CCCTAACTCCGCCCATCCCGCCCCTAACTCCGCCCAGTTCCGCCCATTCTCCGCCC<br>CATGGCTGACTAATTTTTTTTATTTATGCAGAGGCCGAGGCCGCCTCTGCCTCTGA<br>GCTATTCCAGAAGTAGTGAGGAGGCTTTTTTGGAGGCCTAGGCTTTTGCAAAAAGC<br>TCCCGGGAGCTTGTATATCCATTTTCGGATCTGATCAGCACGTGTTGACAATTAAT<br>CATCGGCATAGTATATCGGCATAGTATAATACGACAAGGTGAGGAACTAAACCATG<br>GCCAAGTTGACCAGTGCCGTTCCGGTGCTCACCGCGCGCGACGTCGCCGGAGCGGT<br>CGAGTTCTGGACCGACCGGCTCGGGTTCTCCCGGGACTTCGTGGAGGACGACTTCG<br>CCGGTGTGGTCCGGGACGACGTGACCCTGTTCATCAGCGCGGTCCAGGACCAGGTG<br>GTGCCGGACAACACCCTGGCCTGGGTGTGGGTGCGCGGCCTGGACGAGCTGTACGC<br>CGAGTGGTCGGAGGTCGTGTCCACGAACTTCCGGGACGCCTCCGGGCCGGCCATGA<br>CCGAGATCGGCGAGCAGCCGTGGGGGCGGGAGTTCGCCCTGCGCGACCCGGCCGGC<br>AACTGCGTGCACTTCGTGGCCGAGGAGCAGGACTGACACGTGCTACGAGATTTCGA<br>TTCCACCGCCGCCTTCTATGAAAGGTTGGGCTTCGGAATCGTTTTCCGGGACGCCG<br>GCTGGATGATCCTCCAGCGCGGGGATCTCATGCTGGAGTTCTTCGCCCACCCCAAC<br>TTGTTTATTGCAGCTTATAATGGTTACAAATAAAGCAATAGCATCACAAATTTCAC<br>AAATAAAGCATTTTTTTCACTGCATTCTAGTTGTGGTTTGTCCAAACTCATCAATG<br>TATCTTATCATGTCTGTATACCGTCGACCTCTAGCTAGAGCTTGGCGTAATCATGG<br>TCATAGCTGTTTCCTGTGTGAAATTGTTATCCGCTCACAATTCCACACAACATACG<br>AGCCGGAAGCATAAAGTGTAAAGCCTGGGGTGCCTAATGAGTGAGCTAACTCACAT<br>TAATTGCGTTGCGCTCACTGCCCGCTTTCCAGTCGGGAAACCTGTCGTGCCAGCTG<br>CATTAATGAATCGGCCAACGCGCGGGGAGAGGCGGTTTGCGTATTGGGCGCTCTTC<br>CGCTTCCTCGCTCACTGACTCGCTGCGCTCGGTCGTTCGGCTGCGGCGAGCGGTAT<br>CAGCTCACTCAAAGGCGGTAATACGGTTATCCACAGAATCAGGGGATAACGCAGGA<br>AAGAACATGTGAGCAAAAGGCCAGCAAAAGGCCAGGAACCGTAAAAAGGCCGCGTT<br>GCTGGCGTTTTTCCATAGGCTCCGCCCCCCTGACGAGCATCACAAAAATCGACGCT<br>CAAGTCAGAGGTGGCGAAACCCGACAGGACTATAAAGATACCAGGCGTTTCCCCCT<br>GGAAGCTCCCTCGTGCGCTCTCCTGTTCCGACCCTGCCGCTTACCGGATACCTGTC<br>CGCCTTTCTCCCTTCGGGAAGCGTGGCGCTTTCTCAATGCTCACGCTGTAGGTATC<br>TCAGTTCGGTGTAGGTCGTTCGCTCCAAGCTGGGCTGTGTGCACGAACCCCCCGTT<br>CAGCCCGACCGCTGCGCCTTATCCGGTAACTATCGTCTTGAGTCCAACCCGGTAAG<br>ACACGACTTATCGCCACTGGCAGCAGCCACTGGTAACAGGATTAGCAGAGCGAGGT<br>ATGTAGGCGGTGCTACAGAGTTCTTGAAGTGGTGGCCTAACTACGGCTACACTAGA<br>AGGACAGTATTTGGTATCTGCGCTCTGCTGAAGCCAGTTACCTTCGGAAAAAGAGT<br>TGGTAGCTCTTGATCCGGCAAACAAACCACCGCTGGTAGCGGTGGTTTTTTTGTTT<br>GCAAGCAGCAGATTACGCGCAGAAAAAAAGGATCTCAAGAAGATCCTTTGATCTTT<br>TCTACGGGGTCTGACGCTCAGTGGAACGAAAACTCACGTTAAGGGATTTTGGTCAT |

| | | |
|---|---|---|
| | | GAGATTATCAAAAAGGATCTTCACCTAGATCCTTTTAAATTAAAAATGAAGTTTTA<br>AATCAATCTAAAGTATATATGAGTAAACTTGGTCTGACAGTTACCAATGCTTAATC<br>AGTGAGGCACCTATCTCAGCGATCTGTCTATTTCGTTCATCCATAGTTGCCTGACT<br>CCCCGTCGTGTAGATAACTACGATACGGGAGGGCTTACCATCTGGCCCCAGTGCTG<br>CAATGATACCGCGAGACCCACGCTCACCGGCTCCAGATTTATCAGCAATAAACCAG<br>CCAGCCGGAAGGGCCGAGCGCAGAAGTGGTCCTGCAACTTTATCCGCCTCCATCCA<br>GTCTATTAATTGTTGCCGGGAAGCTAGAGTAAGTAGTTCGCCAGTTAATAGTTTGC<br>GCAACGTTGTTGCCATTGCTACAGGCATCGTGGTGTCACGCTCGTCGTTTGGTATG<br>GCTTCATTCAGCTCCGGTTCCCAACGATCAAGGCGAGTTACATGATCCCCCATGTT<br>GTGCAAAAAAGCGGTTAGCTCCTTCGGTCCTCCGATCGTTGTCAGAAGTAAGTTGG<br>CCGCAGTGTTATCACTCATGGTTATGGCAGCACTGCATAATTCTCTTACTGTCATG<br>CCATCCGTAAGATGCTTTTCTGTGACTGGTGAGTACTCAACCAAGTCATTCTGAGA<br>ATAGTGTATGCGGCGACCGAGTTGCTCTTGCCCGGCGTCAATACGGGATAATACCG<br>CGCCACATAGCAGAACTTTAAAAGTGCTCATCATTGGAAAACGTTCTTCGGGGCGA<br>AAACTCTCAAGGATCTTACCGCTGTTGAGATCCAGTTCGATGTAACCCACTCGTGC<br>ACCCAACTGATCTTCAGCATCTTTTACTTTCACCAGCGTTTCTGGGTGAGCAAAAA<br>CAGGAAGGCAAAATGCCGCAAAAAAGGGAATAAGGGCGACACGGAAATGTTGAATA<br>CTCATACTCTTCCTTTTTCAATATTATTGAAGCATTTATCAGGGTTATTGTCTCAT<br>GAGCGGATACATATTTGAATGTATTTAGAAAAATAAACAAATAGGGGTTCCGCGCA<br>CATTTCCCCGAAAAGTGCCACCTGACGTC |

### SUPPLEMENTARY NOTE

**Thermodynamic modeling of competitive hetero- and homo-dimerization**

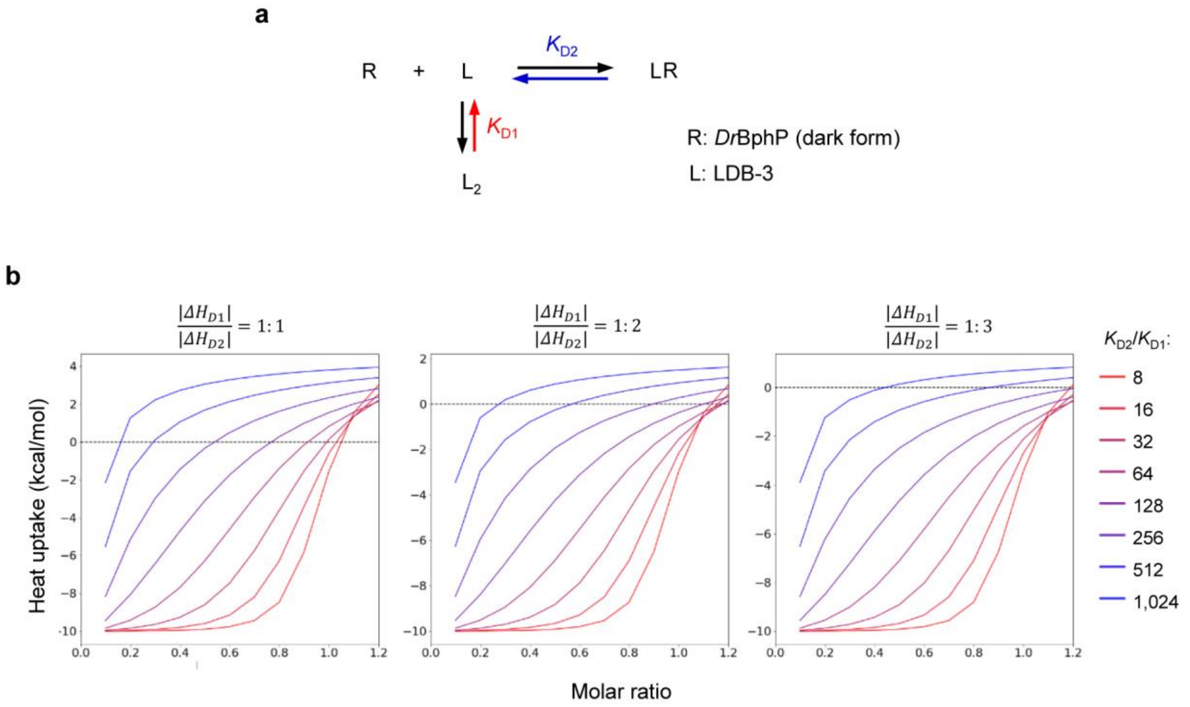

**Figure S12. (a)** Two-state equilibrium model used in the thermodynamic simulation. (**b**) Thermographs showing the integration of heat transfer in simulated titration experiments.

A simplified thermodynamic model was used to understand the observed transition of heat transfer from heat release to absorption when titrating LDB-3 to the dark-form DrBphP (Fig. 5a). We assume that the dark-form *Dr*BphP–LDB-3 binding and LDB-3 dimerization are competitive (Fig. S12a): $2L \leftrightarrow L_2$, and $L + R \leftrightarrow LR$, where L represents the monomeric LDB-3 and R the dark-form *Dr*BphP. The dissociation constants are $K_{\mathrm{D1}} = \frac{[L]^2}{[L_2]} = \frac{1}{K_{a1}}$ and $K_{\mathrm{D2}} = \frac{[L][R]}{[LR]} = \frac{1}{K_{a2}}$, where [L], [$L_2$], [R], and [LR] are equilibrium concentrations, and $K_{a1}$ and $K_{a2}$ are association constants. The relationships of these concentrations are $[L_T] = [L] + [LR] + 2[L_2]$, and $[R_T] = [R] + [LR]$, where [$L_T$] represents the initial total concentration of LDB-3 and [$R_T$] represents the initial total concentration of *Dr*BphP. So, the equilibrium dissociation constants can also be expressed as $K_{D1} = \frac{([L_T]-2[L_2]-[LR])^2}{[L_2]}$, and $K_{D2} = \frac{([L_T]-2[L_2]-[LR])([R_T]-[LR])}{[LR]}$. [$L_2$] and [LR] could be determined if [$K_{D1}$], [$K_{D2}$], [$L_T$] and [$R_T$] are known.

The equilibrium dissociation constant is associated with the Gibbs energy of dissociation, $\Delta G_D$, and can be expressed in terms of the enthalpy($\Delta H_D$) and entropy ($\Delta S_D$) changes in the process: $\Delta G_D = -RT \ln K_D = \Delta H_D - T\Delta S_D$. During the ITC assay, we assume that the whole heat transfer is the sum of $\Delta H_{D1}$ and $\Delta H_{D2}$ which could be calculated by concentration changes of each component using above equations. To simulate a titration process, the dissociation of the LDB-3 homodimer was set to be *endothermic* ($\Delta H_{D1} > 0$) while the formation of the LDB-3-*Dr*BphP complex was *exothermic* ($\Delta H_{D2} < 0$), which is consistent with our experimental results (Figs. 5a and S7). To calculate heat transfer of the whole system, $K_{D2}/K_{D1}$ was set as a variable, and $\frac{|\Delta H_{D1}|}{|\Delta H_{D2}|}$ was set to be 1:1, 1:2, or 1:3. Thermographs were generated to show the integration of heat transfer in an titration experiment. Below is the command used to run the simulation:

```
import os
import os.path
import sys
from scipy.optimize import fsolve
from matplotlib import pyplot as plt

n_point = 13 # number of drops
L2_lst = [] # the concentration of L2 after each drop
LR_lst = [] # the concentration of LR after each drop
R_lst = [] # the concentration of R after each drop
L_lst = [] # the concentration of L after each drop
for i in range(0, n_point):
      ka1 = 1e5 #ka1
      ka2 = 200 #ka2
      R0 = 1 #Rt
      if i == 0:
            L0 = R0 *  1e-6 / 10 #Lt, avoid 0 in calculation
      else:
            L0 = R0 *  i / 10 # Lt
      results  =  solve([Eq(L2-ka1  *  (L0-2*L2-LR)*(L0-2*L2-LR),  0),  Eq(LR-
ka2*(R0-LR)*(L0-2*L2-LR),0)], [L2, LF]) # solve equations
      L2_lst.append(results[0][0].as_real_imag()[0])
      LR_lst.append(results[0][1].as_real_imag()[0])
      R_lst.append(R0-results[0][1].as_real_imag()[0])
      L_lst.append((L0-results[0][1].as_real_imag()[0]-
2*results[0][0].as_real_imag()[0])/2)

delta_L2_lst = [] #the change of concentration of L2 between two drops
delta_LR_lst = [] #the change of concentration of LR between two drops
delta_R_lst = [] #the change of concentration of R between two drops
delta_L_lst = [] #the change of concentration of L between two drops

for i in range(1, len(L2_lst)):
      delta_L2_lst.append(L2_lst[i]-L2_lst[i-1])
      delta_LR_lst.append(LR_lst[i]-LR_lst[i-1])
      delta_R_lst.append(R_lst[i]-RF_lst[i-1])
      delta_L_lst.append(L_lst[i]-L_lst[i-1])
```

```
dLR = -100 #the heat change of L+R->LR
dL2 = 50 # the heat change of L+L->L2
x_lst = list(range(1, n_point)) #molar ratio
for i in range(len(x_lst)):
      x_lst[i] = float(x_lst[i])/10
H_lst = [] #data change
for i in range(len(delta_L2_lst)):
      H_lst.append(dLR * delta_LR_lst[i] + dL2 * delta_L2_lst[i])

# generate plots
plt.figure(figsize=(10,10))
plt.plot(x_lst, H_lst)
plt.xlabel("Molar Ratio")
plt.ylabel("Heat uptake")
```

The simulation result showed that the clear transition from the heat release to absorption was found when $K_{D2}$ >> $K_{D1}$ (e.g., $K_{D2}/K_{D1}$ > 100). The LDB-3 dimer is expected to a relatively weak complex because, in the SEC experiment, a large percentage of the dimer was dissociated at the low-µM concentrations (Figure S6a). Based on our simulation and observed data (Fig. 5a), the dark-form *Dr*BphP–LDB-3 complex ($K_{D2}$) should be much weaker than the LDB-3 dimer ($K_{D1}$), 3. It should be noted that this simplified model did not consider *Dr*BphP dimerization and possible binding cooperativity in higher-order complexes. The fitting of the dark form binding data was found to be difficult due to the complexity of protein-protein interactions, so we did not calculate the $K_D^{app}$. Nevertheless, the ITC experimental data and the modeling supports the low dark activity of LDB-3 observed in other assays.
